# Commensal-specific CD4 T cells promote inflammation in the central nervous system via molecular mimicry

**DOI:** 10.1101/2022.09.27.509786

**Authors:** Zachary White, Ivan Cabrera, Andrea Ochoa-Raya, Isabel Kapustka, Kevin P. Koster, Takashi Matsuo, Dulari Jayawardena, Takahiro Kageyama, Pradeep K Dudeja, Sarah E. Lutz, Akira Yoshii, Teruyuki Sano

**Author notes:** Contributed equally (Zachary White, Ivan Cabrera).

## Abstract

Commensal bacteria are critical regulators of both tissue homeostasis and the development and exacerbation of autoimmunity. However, it remains unclear how the intestinal microbiota contributes to inflammation in tissues such as the central nervous system (CNS) where these microbes are typically absent and whether T cell receptor (TCR) specificity for commensal-derived antigens is important to the development of tissue inflammation-related outcomes. Here, we found that ileum- and cecum-colonizing segmented filamentous bacteria (SFB)-specific T cells (clone TCR^7B8^) can infiltrate the CNS wherein they can be reactivated and produce high levels of inflammatory cytokines including IFNγ, IL-17A, TNFα, and GM-CSF in the absence of regulatory T cells. In contrast, other SFB-specific T cells (clone TCR^1A2^) recognizing an epitope in which 8/9 amino acids overlap with those recognized by TCR^7B8^ failed to induce such neuroinflammation. Despite their similar SFB-derived peptide antigen targets, TCR^7B8^ was found to recognize peptides derived from host proteins including receptor tyrosine-protein kinase ErbB2, trophinin 1, and anaphase-promoting complex subunit 2 *in vitro*, whereas TCR^1A2^ did not, indicating that TCR^7B8^ induces CNS inflammation via molecular mimicry. Immune checkpoint blockade accelerated TCR^7B8^-mediated CNS inflammation, suggesting a potential cause of immune-related adverse events induced in cancer patients undergoing such treatment. Together, our findings reveal a potential mechanism whereby gut commensal-specific T cells are dysregulated and contribute to extraintestinal inflammation.

## Main

The microbiota has been recognized as a critical regulator of various diseases^1,2,3^, with numerous studies having reported changes in the composition of the gut microbiome in the context of not only inflammatory bowel disease (IBD)^4,5,6^, but also neurodegenerative diseases^7,8^. These microbial shifts can exert systemic effects by altering the release of specific metabolites into the bloodstream^9-11^. The gastrointestinal microbiota has further been reported to exhibit immunomodulatory activity capable of activating innate and adaptive immunity^12,13^. While few commensal bacteria have been confirmed to induce antigen-specific T cell responses, commensal-specific T cells are nonetheless thought to provide diverse T cell receptor (TCR) repertoires that supply a variety of T cell pools^14^, particularly given that these commensal-specific T cells can enter systemic circulation^15,16^. Some commensals contribute to not only the generation of protective and homeostatic immunity but also the onset and severity of several autoimmune diseases^17^. However, it remains unclear as to whether the specificity of TCRs for commensal-derived antigens is important to the development of tissue inflammation.

Segmented filamentous bacteria (SFB) are ileum- and cecum-colonizing spore-forming gram-positive commensal bacteria^18,19^ that have been detected in humans^20,21^ and other vertebrates, as well as in invertebrates^22,23^. SFB are immunogenic and can induce the development of Th17 cells specific for SFB-derived antigens in the murine intestines^24, 25, 26^. While SFB predominantly thrive in the terminal ileum and cecum^27^, SFB colonization in various murine autoimmune disease models results in the inflammation of the joints and several other distal tissues^28,29^. Three SFB-specific TCR transgenic (TCR Tg) mouse lines have been established to date^26^. When naïve CD4 T cells are adaptively transferred from these SFB TCR Tg mice into wild-type (WT) C57BL/6 hosts, they differentiate into Th17 cells in the gut-draining mesenteric lymph nodes (mLNs) in response to SFB^30^. A majority of these expanded SFB-specific CD4 T cells are maintained in the intestines where the commensals they recognize are located^16^, yet a subset of these cells are detectable in distal tissues including the spleen and lungs^16,31^. As such, SFB TCR Tg mice represent an ideal tool for use when exploring the relationships between commensal-derived antigen specificity and the pathogenesis of tissue inflammation.

### SFB-specific TCR^7B8^ induces inflammation in the intestines and central nervous system in immunodeficient hosts

To elucidate the requirement for SFB-specific TCR recognition as a driver of inflammation in the gut and other tissues, we utilized a well-characterized transfer colitis model system^32,33^. In this system, in the absence of regulatory T cells (Tregs)^34,35^, the transfer of bulk naïve CD4 T cells (carrying diverse heterogeneous TCRs) from WT mice into immunocompromised hosts results in the differentiation of these cells into pathogenic CD4 T cells, inducing colitis in a commensal-dependent manner^36,37^. To fix the TCR on naïve T cells in this model, we leveraged SFB-specific TCR transgenic (SFB TCR^7B8^ Tg) mice as T cell donors. After we transferred 1×10^5^ naïve CD4 T cells from SFB TCR^7B8^ Tg mice into SFB-colonized *Rag2*^*-/-*^ hosts, the recipient animals exhibited symptoms of wasting (Fig. 1a) and inflammation in the ileum and colon (Fig. 1b, c). Unexpectedly, these recipient mice also developed a hind-limb clasping phenotype (Fig. 1d, e, Supplemental Video 1-3 and Extended Data Fig. 1a) that affected 100% of these mice (Fig. 1f and Extended Data Fig. 1b). Some transferred mice exhibited very severe clasping of not only their hind legs, but also their forelegs (Supplemental Video 4). Unlike mice into which myelin-specific naïve T cells (TCR^2D2^) have been transferred, which develop experimental autoimmune encephalomyelitis (EAE) characterized by the paralysis of both the legs and tail^38^, the tails of SFB TCR^7B8^-recipient mice were completely normal, and limb clasping was observed to similar extents in both males and females (Fig. 1e, f). In healthy mice, few or no CD4 T cells were generally detectable in the brain or spinal cord, whereas the number of IFNγ-producing or Tbet-positive Th1 cells detected in the CNS rose markedly in TCR^7B8^-transferred recipient mice (Fig. 1g-j, Extended Data Fig. 1c and 2). Consistent with the observed intestinal pathology, we detected IFNγ-producing Th1, IL-17A-producing Th17, and IFNγ- and IL-17A-double producing (DP) pathogenic T cells^39,40^ in the ileum and colon of these mice (Fig. 1i). In addition, substantially more IFNγ/IL-17A DP and IFNγ/TNFα DP T cells were detected in the brain and spinal cord of these animals (Fig. 1i-k).

**Fig. 1.**
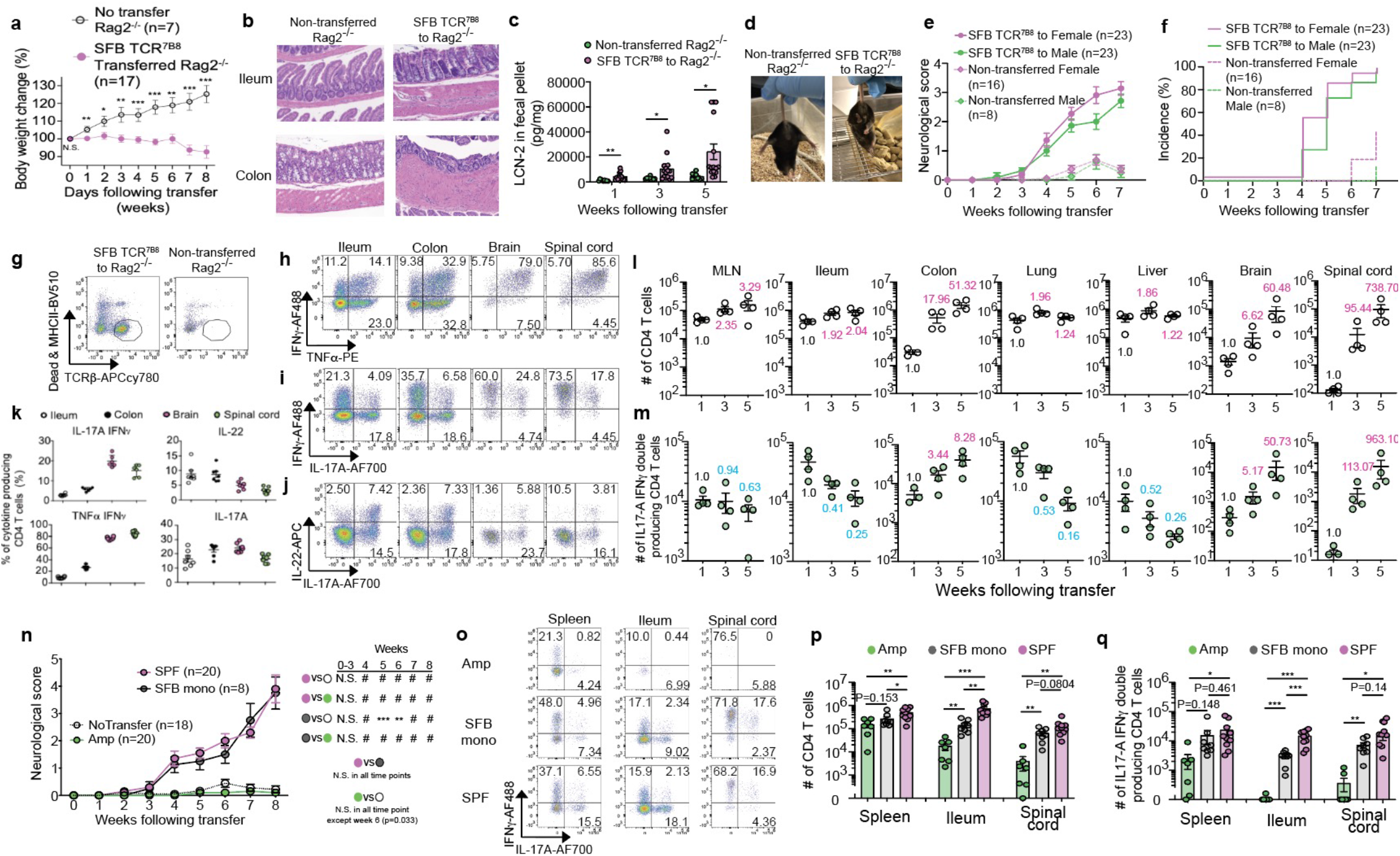
Transferred SFB TCR^7B8^ CD4 T cells expand and exhibit pathogenic activity in the intestines an SFB antigen-dependent manner. **a-m**, 1 × 10^5^ SFB TCR^7B8^ CD4 naïve T cells were retro-orbitally transferred into SFB-colonized *Rag2*^*-/-*^ hosts. After transfer, animals were assessed for changes in t (**a**), tissue pathology as analyzed via H&E staining (n=5/group) (**b**), and LCN2 levels in freshly cal pellets at the indicated weeks following the transfer is measured by ELISA (n=9 non-*Rag2-/-* and n=15 SFB TCR^7B8^ transferred *Rag2*^*-/-*^ mice). (**c**), representative images of hind limb henotypes (**d**), hind limb clasping scores (**e**), and hind limb clasping incidence (**f**). Four t experiments with multiple replicates comprised primarily of littermates were conducted, and all ta was combined. n=23 (transferred *Rag2*^*-/-*^ female mice, transferred *Rag2*^*-/-*^ male mice), n=16 erred *Rag2*^*-/-*^ female mice), n=8 (non-transferred Rag2^-/-^ male mice). Representative FACS plots T cells (**g**), IFNγ- and TNFαξ-(**h**), IL-17A-(**i**), and IL-22-producing (**j**) CD4 T cells in the ileum,, and spinal cord of transferred *Rag2*^*-/-*^ hosts at 8 weeks post-transfer. **k**, The frequency of oducing cells among CD4 T cells in transferred *Rag2*^*-/-*^ hosts 8 weeks post-transfer. Data were rom two independent experiments (n=6). **l, m**, TCR^7B8^ CD4 T cell kinetics in the mLN, ileum, liver, brain, and spinal cord of Rag2^-/-^ hosts at 1-, 3-, and 5-weeks post-transfer. Total CD4 (**l**) -17A double-producing CD4 T cell counts (**m**); four biological replicates per time point. **n-q**, eated or SFB mono-colonized *Rag2*^*-/-*^ hosts received a retro-orbital transfer of 1 × 10^5^ SFB 4 naïve T cells. Neurological disorder scores were measured from two experiments (**h**), n=20 to SFB+ *Rag2*^*-/-*^ hosts under SPF conditions), n=8 (SFB mono-colonized *Rag2*^*-/-*^ mice), n=20 d SFB-*Rag2*^*-/-*^ hosts), n=20 (Non-transferred Rag2^-/-^ hosts). Representative FACS plots ing IFNγ and IL-17A expression (**o**), total CD4 T cell counts (**p**), and IFNγ IL-17A double D4 T cell counts (**q**) in the spleen, ileum, and spinal cord from the transferred *Rag2*^*-/-*^ hosts at 8 -transfer. Two independent experiments were combined. n=7 (Amp), n=7 (SFB mono), n=10 ent’s t-test. #P<0.0001, ***P<0.001, **P<0.01, N.S. not significant (p>0.05).

Using this system, we next sought to understand when, where, and how the pathogenic status of T cells in various tissues is induced, maintained, or altered following adoptive T cell transfer. We collected mononuclear cells from the lungs, liver, mLNs, ileum, colon, brain, and spinal cord at 1, 3, and 5 weeks following SFB TCR^7B8^ naïve CD4 T cell transfer and analyzed effector cytokine production. At 1-week post-transfer, SFB TCR^7B8^ CD4 T cells were not detected in the brain or spinal cord, whereas IL-17A/IFNγ-DP TCR^7B8^ CD4 T cells were present not only in the mLNs, ileum, and colon wherein SFB antigens are generally abundant, but also in the lungs and liver (Fig. 1l, m). Importantly, these pathogenic T cells did not exhibit long-term retention in the liver or lungs, where SFB are generally absent (Fig. 1m). Although SFB-derived antigens should be abundant in the ileum, the observed ileal pathogenic T cell population also contracted over time but was maintained to some extent. At later time points, pathogenic CD4 T cell numbers increased in the colon, spinal cord, and brain. These results suggested that wasting and hind-limb clasping were caused by chronic tissue inflammation in the colon and CNS in response to SFB-specific naïve T cell transfer, but not by the acute tissue inflammation observed in the lungs and liver. In some cases, T cells from TCR Tg mice express both endogenous and Tg TCRs^41^. While the frequency of these “Dual-TCR”-positive T cells is not high, they nonetheless contribute to the induction of tissue inflammation given that they undergo selective expansion owing to their dual-specificity^31,42^. To elucidate whether leg clasping in our model system was induced via Dual-TCR-related effects, we crossed SFB TCR^7B8^ Tg mice with *Rag2*^*-/-*^ mice. Due to the lack of Rag2, endogenous TCRs cannot undergo TCR rearrangement such that only the Tg TCR is expressed on T cells in TCR Tg *Rag2*^*-*^ */-* mice. Following the transfer of naïve SFB TCR^7B8^ Tg *Rag2*^*-/-*^ T cells into *Rag2*^*-/-*^ hosts, these mice developed similar levels of hind-limb clasping (Extended Data Fig. 3a). Importantly, SFB TCR^7B8^ mice on the *Rag2*^*-/-*^ background developed spontaneous hind-limb clasping without any transfer or treatment in an SFB+ specific pathogen-free (SPF) facility (Extended Data Fig. 3b). Moreover, SFB TCR^7B8^ *Rag2+/-* mice developed milder clasping than mice on the *Rag2*^*-/-*^ background at later ages owing to the presence of Treg cells (Extended Data Fig. 3b-c).

### SFB colonization is necessary and sufficient to induce TCR^7B8^-mediated CNS inflammation

Next, to establish whether SFB colonization is necessary for the development of CNS inflammation, we treated SFB-colonized *Rag2*^*-/-*^ hosts with Ampicillin (Amp) to preventatively remove SFB prior to T cell transfer^43^ (Extended Data Fig. 4a). Mice treated with Amp before the transfer did not develop any wasting (Extended Data Fig. 4b) or leg clasping (Fig. 1n). Owing to the homeostatic proliferation of transferred naïve T cells in these immunocompromised hosts^44,45^, some transferred CD4 T cells in the spleen of Amp-treated SFB negative hosts were found to have expanded in an antigen-independent fashion, but these cells did not produce high levels of effector cytokines (Fig. 1o-q). Moreover, SFB TCR^7B8^ CD4 T cells expanded and activated by homeostatic proliferation were not detected in the spinal cord, and the recipient hosts did not develop clasping phenotypes (Fig. 1n, o). To further elucidate whether the SFB colonization of these animals is sufficient to induce clasping phenotypes, we transferred SFB TCR^7B8^ naïve T cells into *Rag2*^*-/-*^ hosts mono-colonized with SFB inside a germ-free (GF) isolator to prevent contamination with any other commensal species during the entirety of the experiment (Extended Data Fig. 4c). Clasping was still observed in these SFB mono-colonized hosts (Fig. 1n) and activated SFB TCR^7B8^ CD4 T cells were present in their spinal cords, in line with what was observed in SFB+ *Rag2*^*-/-*^ mice under SPF conditions (Fig. 1o-q). Taken together, these results led us to conclude that the infiltration, expansion, and dysregulation of SFB-TCR^7B8^ T cells in the CNS was initiated by SFB-derived antigens, while their pathogenicity is likely attributable to local environmental factors within the CNS.

### TCR^7B8^ CD4 T cells produce GM-CSF and activate microglia in the CNS

To understand how infiltrating SFB TCR^7B8^ CD4 T cells contribute to the development of clasping phenotypes, we hypothesized that these cells may produce a range of inflammatory cytokines and accumulate in specific locations wherein motor neurons are present and/or project in the CNS. Consistently, infiltrating SFB TCR^7B8^ CD4 T cells in the brain and spinal cord produced more IFNγ and GM-CSF than in the ileum or colon (Fig. 2a, b). We therefore surveyed activated microglial localization within the diseased brain tissue of TCR^7B8^ T cell-transferred *Rag2*^*-/-*^ hosts. Relative to non-diseased hosts, we detected more activated microglia, as defined by Iba-1 positivity^46^ and ameboid microglial morphology^47^, in the brainstem, cerebrum, motor cortex, striatum, and thalamus wherein motor neurons are located^48^ (Fig. 2c). In a myelin oligodendrocyte glycoprotein, (MOG)-immunization-induced EAE model, CD4 T cells were observed in the myelin-enriched white matter of the spinal cord^49^, whereas we detected SFB TCR^7B8^ CD4 T cells in the spinal cord grey matter (Fig. 2d-e). Consistent with this observation, myelin located in the white matter of the spinal cord in the TCR^7B8^-transferred hosts was intact (Extended Data Fig. 5). These results suggest that TCR^7B8^ CD4 T cells may activate microglia by producing GM-CSF and contribute to CNS inflammation through mechanisms distinct from those observed in EAE.

**Fig. 2.**
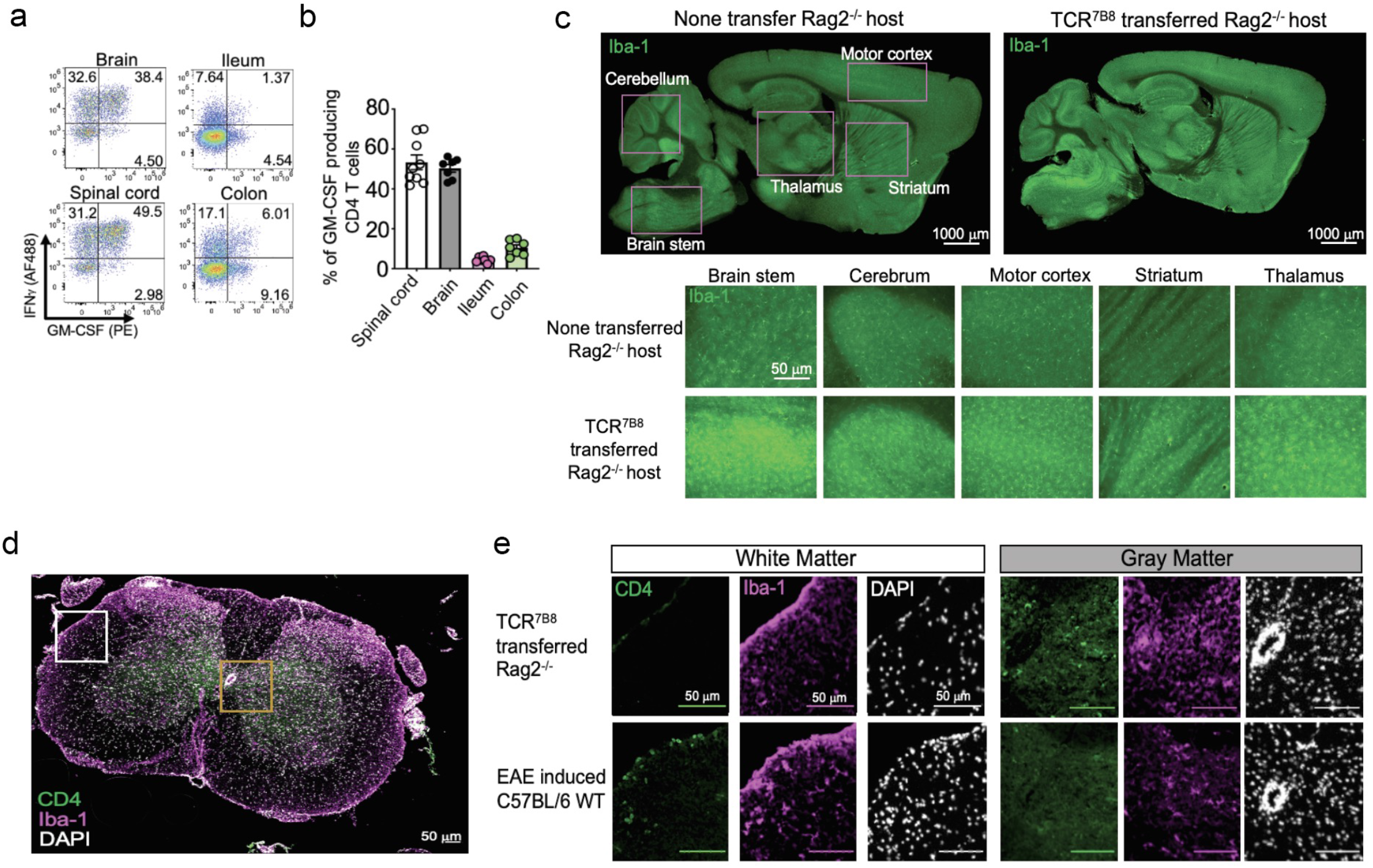
Pathogenic SFB TCR^7B8^ CD4 T cells induce site-specific inflammation. **a-e**, SFB-colonized s received a retro-orbital transfer of 1 × 10^5^ SFB TCR^7B8^ CD4 naïve T cells. IFNγ and GM-CSF in CD4 T cells from the spinal cord, brain, ileum, and colon of these *Rag2*^*-/-*^ hosts was assessed post-transfer. Flow plots of IFNγ and/or GM-CSF expressing CD4 T cells (**a**) and the percentage -producing cells among CD4 T cells (**b**). Two independent experiments were combined (n=9). **c**, tive images of brain sections from TCR^7B8^ CD4 T cell recipient *Rag2*^*-/-*^ hosts at 8 weeks post-mples were stained for Iba-1 (green). **d**, Representative images of spinal cord sections from d WT mice (**e**, bottom), and *Rag2*^*-/-*^ hosts 8 weeks post-transfer of SFB TCR^7B8^ naïve T cells (**e**, d for Iba-1 (magenta), CD4 (green), and DAPI (white). White and orange rectangles shown in **d** ed in **e**. This experiment was repeated twice with similar staining results.

### Non-TCR^7B8^ SFB-specific TCR expression is insufficient to mediate CD4 T cell-induced CNS inflammation

We next evaluated how SFB TCR^7B8^ T cells can infiltrate, accumulate, and/or be re-activated in the CNS. To determine whether SFB TCR^7B8^ CD4 T cells undergo active proliferation within the CNS, we analyzed the expression of the proliferation marker Ki67 on transferred T cells in various tissues. To mitigate the influence of antigen-independent homeostatic proliferation, we focused on the chronic spontaneous clasping phenotypes observed in SFB TCR^7B8^ *Rag2*^*-/-*^ mice rather than in adoptively transferred animals (Extended Data Fig. 3b). CD4 T cells in the spinal cord, brain, ileum, and colon from SFB TCR^7B8^ *Rag2*^*-/-*^ mice expressed Ki67, although those in the spleen did not (Extended Data Fig. 6), suggesting that SFB TCR^7B8^ CD4 T cells were actively stimulated and proliferating in the CNS. However, SFB was not detectable in the CNS in these animals (data not shown). When recipients of TCR^7B8^ naïve T cells were treated with Amp following the development of neurological phenotypes to remove SFB from these diseases animals, their neurological scores failed to decrease despite the elimination of SFB-derived antigens (Extended Data Fig. 7). This suggested that while SFB-derived antigens are required for the expansion and dysregulation of SFB-specific TCR^7B8^ T cells in the gut, these SFB-specific TCR^7B8^ T cells may be restimulated in an SFB-independent manner within the CNS following initial expansion in the gastrointestinal compartment and systemic dissemination. Therefore, we hypothesized that SFB TCR^7B8^ T cells may be stimulated by MHCII molecules presenting cross-reactive host peptides within the CNS. To test this possibility, we prepared another SFB-specific TCR Tg model, SFB TCR^1A2^ mice, which generate Tg T cells that recognize the same SFB-derived protein, SFB_003340, with similar affinity to that of SFB TCR^7B8 26^. Much like TCR^7B8^, TCR^1A2^ is one of the dominant SFB-specific TCR clones in characterized T cell repertoires, yet the amino acid sequences of their TCR α and β β chains are completely different^26^. As shown in Extended Data Fig. 3b, SFB TCR^7B8^ *Rag2*^*-/-*^ mice spontaneously developed clasping phenotypes at 8 weeks of age, and this clasping grew progressively worse over time (Fig. 3a). Consistent with the transient naïve T cell transfer model utilized in Fig. 1, more CD4 T cells and IFNγ/IL-17A DP CD4 T cells were detectable in the brain and spinal cord of SFB TCR^7B8^ *Rag2*^*-/-*^ mice compared with non-TCR Tg *Rag2+/-* littermates (Fig. 3b-d). In contrast, SFB TCR^1A2^ *Rag2*^*-/-*^ mice did not exhibit these phenotypes or corresponding pathological changes (Fig. 3e-h). These results were also confirmed via the transfer of SFB TCR^7B8^ or TCR^1A2^ naïve CD4 T cells into *Rag2*^*-/-*^ hosts (Extended Data Fig. 8). These data indicate that despite their shared specificity for SFB, TCR^7B8^ CD4 T cells but not TCR^1A2^ CD4 T cells undergo active stimulation within the CNS, presumably due to the presence of cross-reactive host antigens in this compartment.

**Fig. 3.**
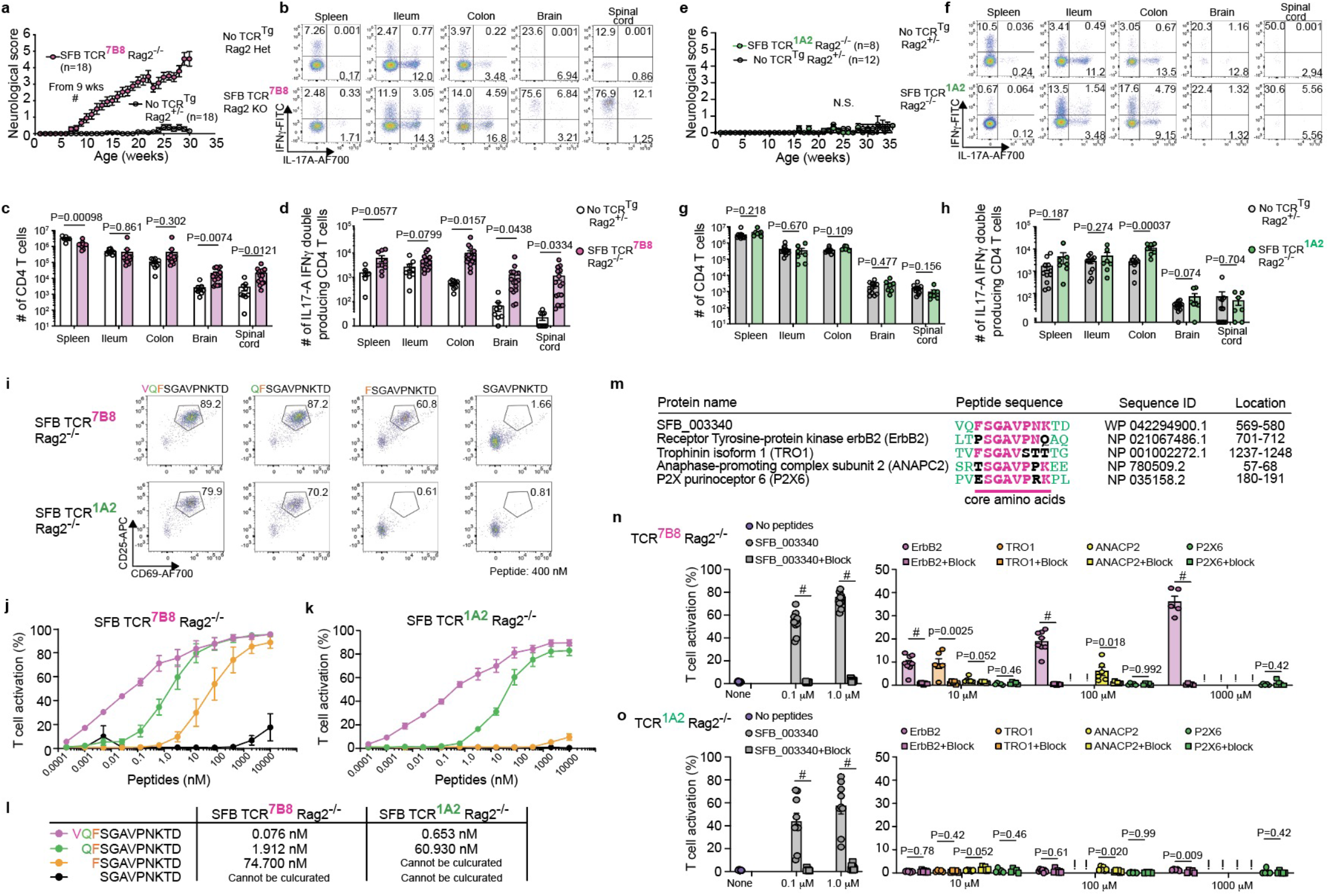
SFB TCR^7B8^ but not SFB TCR^1A2^ contributes to CNS inflammation via the recognition of the cross-st proteins ErbB2, TRO, and ANAPC2. **a-d**, Neurological phenotypes in SFB TCR^7B8^ *Rag2*^*-/-*^ mice R Tg *Rag2+/-* littermate controls. **a**, Neurological scores. All results obtained from multiple littermates ned, and the number of mice per genotype is indicated on the graph. **b-d**, Flow plots corresponding IL-17A expression in CD4 T cells from the indicated tissues of SFB TCR^7B8^ *Rag2*^*-/-*^ mice (**b**), total unts from the indicated tissues in SFB TCR^7B8^ *Rag2*^*-/-*^ mice (**c**), and IFNγ/IL-17A double-producing counts in the indicated tissues of SFB TCR^7B8^ *Rag2*^*-/-*^ mice (**d**). **e-h**, Neurological phenotypes in SFB *2*^*-/-*^ mice compared with non-TCR Tg *Rag2+/-* littermate controls. **e**, Neurological scores. **f-h**, Flow γ and IL-17A expression in CD4 T cells from the indicated tissues of SFB TCR^7B8^ *Rag2*^*-/-*^ mice (**f**), ell counts from the indicated tissues of SFB TCR^1A2^ *Rag2*^*-/-*^ mice (**g**), and IFNγ/IL-17A double-D4 T cell counts from the indicated tissues of SFB TCR^1A2^ *Rag2*^*-/-*^ mice (**h**). All results obtained from rmates were combined, and the number of mice per genotype is indicated on the graph or each dot ented individual mouse. **i-l**. An *in vitro* TCR stimulation assay was performed using bone marrow-dritic cells (*CD45*.*1/CD45*.*1*), naïve T cells from TCR^Tg^ *Rag2*^*-/-*^ mice (*CD45*.*1/CD45*.*2*), and purified rived from the SFB-derived protein, SFB_003340. Flow plots corresponding to CD25 and CD69 in SFB TCR^7B8^ or TCR^1A2^ CD4 T cells (**i**). Kinetics of the TCR^7B8^ (n=3) (**j**) and TCR^1A2^ (n=4) (**k**) n response to truncated peptides side by side. Calculated EC50 values for SFB TCR^7B8^ and TCR^1A2^ o acid alignment. The core epitope for TCR^7B8^ is shown in bold. Shared amino acids are represented **n, o**. An *in vitro* TCR stimulation assay was performed using bone marrow-derived dendritic cells *45*.*1*), naïve T cells from TCR^Tg^ *Rag2*^*-/-*^ mouse (*CD45*.*1/CD45*.*2* or *CD45*.*2/CD45*.*2*), and purified the indicated concentrations, with or without anti-MHCII antibodies. T cell activation was calculated e frequency of CD25 and CD69 expression 24 hours following the co-culturing BMDCs and naïve. !: due to the cell toxity, any co-cultured cells were recovered. Student’s T test. #P < 0.0001, and nificant, or P values were shown in the graph.

It has been reported that the epitope recognized by TCR^7B8^ partially overlaps with that recognized by TCR^1A2 26^. To fully understand the differences in the epitopes recognized by TCR^7B8^ and TCR^1A2^, we prepared the “VGFSGAVPNKTD” peptide, which corresponds to a portion of the SFB_00340 protein present in the murine SFB genome, and assessed its ability to stimulate T cell activation. Bone marrow-derived dendritic cells (BMDCs) were treated with different concentrations of this peptide for 24 hours and then co-cultured with magnetically sorted TCR^7B8^ or TCR^1A2^ Rag2^-/-^ naïve CD4 T cells for an additional 24 hours (Extended Data Fig. 9a). Naïve T cell activation was measured based upon CD25 and CD69 expression (Extended Data Fig. 9b) and also confirmed via Ki67 staining and the dilution of a cell proliferation dye (carboxyfluorescein succinimidyl ester (CFSE)), after 72 hours (Extended Data Fig. 9c). The difference in the affinity of TCR^7B8^ and TCR^1A2^ for this SFB peptide alone can thus not provide a clear explanation as to why TCR^1A2^ Tg mice failed to develop CNS inflammation (Fig. 3j-i). Next, to determine the minimum epitope recognized by both TCR^7B8^ and TCR^1A2^, we utilized a series of truncated peptides for *in vitro* assays. While TCR^7B8^ recognized “FSGAVPNKTD”, TCR^1A2^ failed to respond to this peptide (Fig. 3i-l). This suggests that the 1-2 amino acid difference between the target epitopes of TCR^7B8^ and TCR^1A2^ may belie the specific TCR^7B8^ cross-reactivity observed in our model system.

Next, we conducted a BLAST search for candidate cross-reactive proteins harboring peptide sequences similar to the defined minimum TCR^7B8^ epitope, with 76 candidate peptides identified via this approach then being synthesized to produce a crude peptide library (Extended Fig. 10a). We found that 4 out of these 76 candidate peptides derived from the receptor tyrosine-protein kinase ErbB2, Trophinin 1 (TRO1), anaphase-promoting complex subunit 2 (ANAPC2), and P2X purinoceptor 6 (P2X6) proteins stimulated TCR^7B8^ naïve CD4 T cell activation at 100 μM and 10 μM concentrations (Fig. 3m, Extended Data Fig. 10a-c). These candidates are highly expressed in the CNS according to the Allen Brain Atlas. Confirmatory experiments using synthesized peptides (95% pure) confirmed that ErbB2-, TRO1-, and ANACP2-derived peptides, but not P2X6-derived peptides, were able to stimulate TCR^7B8^ in an MHCII-dependent manner (Fig. 3n). Moreover, none of the tested candidate peptides for TCR^7B8^ did not stimulate TCR^1A2^ naïve CD4 T cells (Fig. 3o). These results indicated that TCR^7B8^ can recognize not only SFB-derived peptides but also at least three murine host proteins expressed in the CNS, potentially contributing to the induction of immunopathological neuroinflammation.

Immune checkpoint molecules expressed on T cells, including CTLA4 and PD-1, play a critical role in preventing inappropriate or excessive effector T cell activation^50,51^. We therefore employed blocking antibodies to attempt to disrupt both PD-1 and CTLA4 signaling in SFB TCR^7B8^ naïve CD4 T cells that had been transferred into SFB-positive *Rag2*^*-/-*^ hosts. As shown in Figure 4, intraperitoneal anti-CTLA4 and anti-PD-1 administration in *Rag2*^*-/-*^ hosts that had undergone T cell transfer resulted in the more rapid development of CNS inflammation that was more severe than that observed in transferred hosts injected with control Abs (Fig. 4a, b). Consistent with the observed disease scores and incidence, checkpoint inhibitor (CPI)-injected hosts also exhibited significantly more total CD4 T cells and activated cytokine-producing CD4 T cells in their CNS as compared with control Ab-injected hosts (Fig. 4c-r). Although CPI-injected TCR^7B8^ transferred hosts presented with far more severe wasting phenotypes (Fig. 4s), intestinal inflammation was comparable between CPI-and control Ab-injected hosts (Fig. 4t), suggesting that the TCR^7B8^ CD4 T cells may undergo further dysregulation during systemic circulation or within the CNS rather than in the intestines in response to the CPI-treatment.

**Fig. 4.**
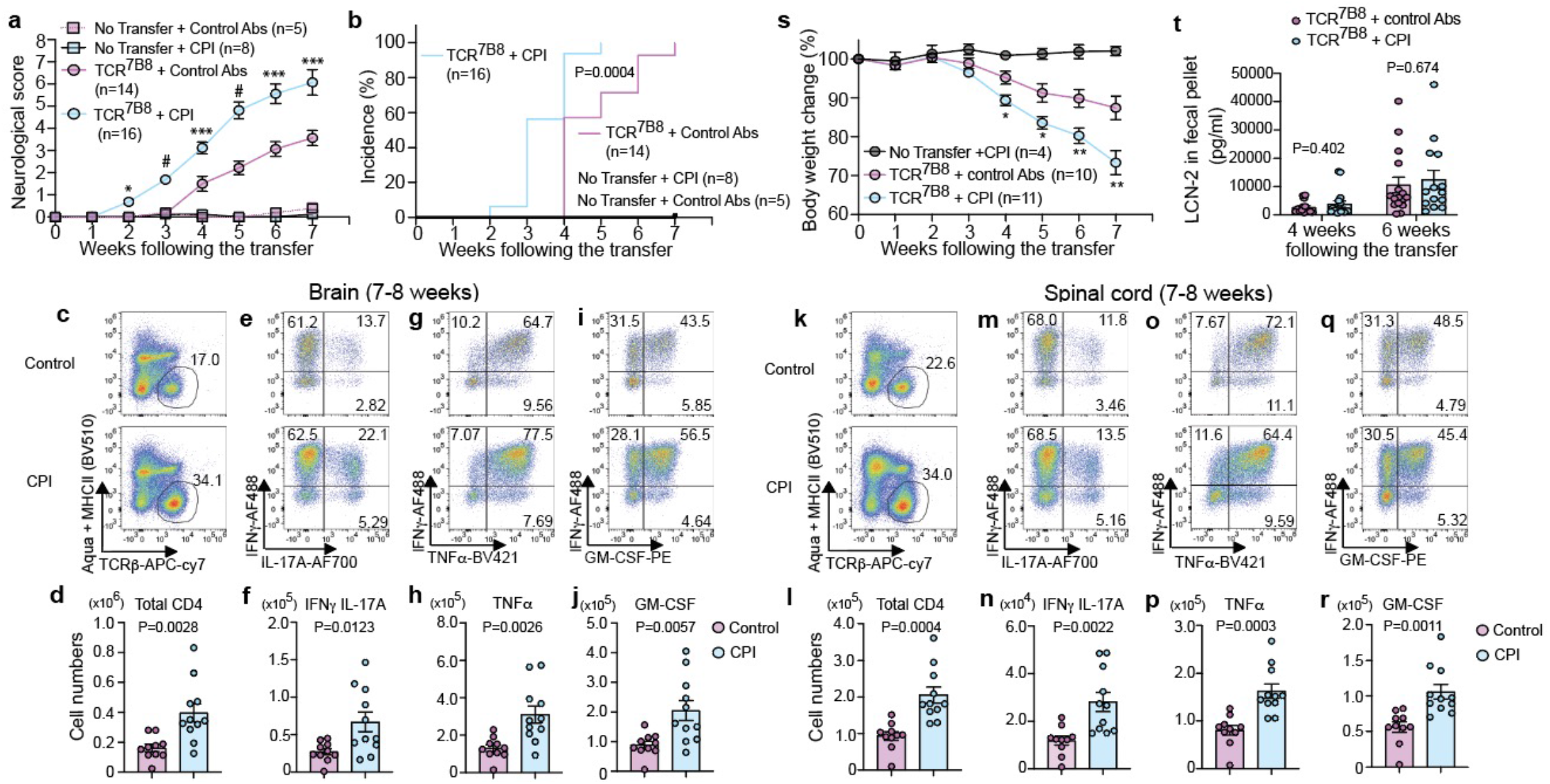
Immune checkpoint blockade promotes TCR^7B8^-mediated CNS inflammation. **Neurological** and incidence (**b**) of the neurological disorders following the transfer of naïve TCR^7B8^ CD4 T cells hosts with or without the i.p. injection of 300 μg of Anti-CTLA4 and Anti-PD-1 antibodies. tive FACS plots and numbers of CD4 T cells in the brain (**c**,**d**) and spinal cord (**k**,**l**), IFNγ/IL-17A ucing CD4 T cells in the brain (**e**,**f**) and spinal cord (**m**,**n**), TNFα-producing CD4 T cells in the and spinal cord (**o**,**p**), GM-CSF-producing CD4 T cells in the brain (**I**,**j**) and spinal cord (**q**,**r**). **s**, t changes over time. **t**, LCN2 detection in freshly collected fecal pellets at the indicated time transfer as measured via ELISA. All the experiments were conducted twice, and the data were ith the exception of the body weight data shown in **s**. *P < 0.05, **P < 0.01, ***P < 0.001, #P < d N.S. Not Significant, or P values were shown in the graph. Student’s T test between TCR^7B8^ hosts injected with control antibodies and TCR^7B8^ transferred hosts injected with CPI.

## Discussion

The above data reveal that SFB-specific T cells are initially generated in the mLNs and intestinal LP in response to SFB-derived antigens, with expanded SFB TCR^7B8^ CD4 T cells subsequently being reactivated in response to peptides derived from host proteins in the CNS via molecular mimicry, resulting in neuroinflammation. While SFB TCR^1A2^ T cells are also generated in the intestines in response to SFB-derived antigens, they fail to undergo restimulation in the CNS, likely owing to the absence of available cross-reactive host peptides. As shown above, the EC50 value corresponding to the interaction between TCR^7B8^ and SFB-derived peptides was very low, consistent with a high-affinity interaction between this TCR and this peptide in the context of its MHCII-dependent presentation. While T cells expressing commensal-specific TCRs can be generated in the thymus^52^, they cannot be eliminated through negative thymic selection, given that commensal genes are not encoded in the host genome or presented by AIRE-expressing thymic epithelial cells. While our data suggest that TCR^7B8^ can additionally recognize host protein-derived peptides, the EC50 value corresponding to the interaction between TCR^7B8^ and these cross-reactive host-derived peptides was much greater than that for SFB-derived peptides. Consequently, these commensal-specific T cells may be able to survive thymic negative selection owing to their low-affinity self-recognition. Our findings, however, highlight unexpected, novel cross-reactivity between commensal-specific TCRs and host proteins (Extended Data Fig. 11), shedding new light on the mechanisms underlying commensal-induced autoimmunity.

Despite the positive clinical impact of the advent of immune CPI treatment in a range of cancer types, in a key trial, 68.7% of patients administered CPIs developed treatment-induced immune-related adverse events (irAEs) characterized by the autoimmune-like inflammation of several organs^53^. Neurological irAEs such as hypophysitis develop in ∼14% of patients after anti-CTLA4 and PD-1 therapy^54^. Although the causes of these irAEs are not understood, commensals are likely to play a role in this process given that they are highly associated with many autoimmune diseases^55,56^. It is thus possible that patients with irAEs harbor commensal-specific T cells that are also cross-reactive to host proteins, with these cross-reactive T cells becoming further dysregulated in response to CPI treatment. Consistent with such a model, gut T cells have previously been shown to contribute to the incidence of CNS inflammation in the context of celiac disease and IBD^57,58^. While celiac disease is caused by the generation of food-reactive T cells and other immune populations, and affected patients experience abdominal pain in the small intestine where food-derived antigens are abundant^59^, a subset of celiac disease patients also exhibit inflammation in the brain^60,61^, presumably due to the migration of dysregulated T cells from the gut to the CNS. Understanding how gut-derived T cells migrate into the brain is critical to developing new treatments capable of preventing T cell-mediated neuroinflammation.

Here, we identified ErbB2, TRO1, and ANAPC2 as cross-reactive proteins recognized by TCR^7B8^ CD4 T cells. These proteins are highly expressed in the brain and spinal cord, but are also expressed in other tissues. While TCR^7B8^ CD4 T cells induce severe inflammation within the CNS, it remains unclear as to whether these cells can induce pathological inflammation in other tissues expressing these cross-reactive host proteins or why this inflammation was restricted to the CNS in the present study. There are two potential explanations for these findings. First, TCR^7B8^ may cross-react not only with ErbB2, TRO1, and ANAPC2, but also with other proteins that are highly abundant in the

CNS that have yet to be identified. Indeed, our crude peptide library experiments revealed pseudo-positive results for the IFT46 and P2X6 proteins when assessing TCR^7B8^ cross-reactivity (Extended Data Fig. 10b, c). Moreover, it is also possible that structurally similar peptides consisting of completely distinct amino acid sequences can be recognized by TCR^7B8 62^.As all possible candidate sequences have not been tested using purified peptides to date, it remains possible that there may be other cross-reactive host proteins within the CNS that are recognized by this TCR. Kishnamoorthy et al. demonstrated that the MOG-specific TCR^2D2^ recognizes not only a MOG-derived peptide but also neurofilament-M (NF-M). TCR^2D2^ T cell transfer into *MOG*^*-/-*^ hosts resulted in delayed EAE, while TCR^2D2^ T cell transfer into *MOG*^*-/-*^ *NF-M*^*-/-*^ hosts resulted in an absence of EAE disease phenotypes^63^. While at least three host proteins may provide autoreactive antigens for TCR^7B8^ CD4 T cells, potentially with some level of redundancy, whether the mutation of these cross-reactive epitopes can alleviate TCR^7B8^-mediated CNS inflammation is worthy of future study.

A second possible explanation for the above results is that the inflammatory cytokines readily produced by TCR^7B8^ CD4 T cells primarily cause inflammation within specific locations such as the brain and spinal cord. In potential support of such a model, Spath et al. previously reported that the dysregulation of GM-CSF induces spontaneous immunopathology in the CNS^64^. Ectopically induced GM-CSF produced by CD4 T cells in an antigen-independent manner can promote spontaneous phagocyte invasion. In the case of TCR^7B8^-mediated CNS inflammation, TCR^7B8^ T cells proliferated, accumulated, and produced GM-CSF within the CNS, but not in the spleen, lung, or liver. Understanding the mechanisms underlying the tissue-restricted nature of these disease-related phenotypes will be of value for efforts to control both tissue-restricted and systemic autoimmune diseases.

The vertebrate gastrointestinal tract is colonized by thousands of microbial species including commensal bacteria, viruses, and fungi, outnumbering the total number of host cells and harboring a combined genomic length substantially longer than that of the host genome^65^. Potential cross-reactive commensal recognition may thus be a cryptic autoimmune risk factor, as was recently reported with respect to the relationship between Epstein-Barr virus-derived protein cross-reactivity and multiple sclerosis^56,66^. Further research exploring the etiology and ubiquity of these commensal-related autoimmune liabilities is thus warranted to inform preventative care and treatment efforts.

## Acknowledgments

We thank Flow Cytometry Core in the UIC Research Resources Center (RRC) for naïve T cell sorting, and Histology core in the RRC for paraffin section. This work was supported by the UIC Dept. of Microbiology and Immunology Startup fund (TS) and Scheweppe Award in Translational Research (TS).

## Author contributions

T.S. designed, performed most experiments, and analyzed the data. Z.W. and I.C. performed most of the experiments. A.O., I.K., T.K. helped with certain experiments. K.P.K., A.O., and T.M. conducted IHC staining for brain and spinal cord samples under the supervision of A.Y., D.J., and P.K.D. contributed the analysis of gut pathology, and

S.E.L. provided spinal cords from EAE-induced mice. T.S. and Z.W. wrote the manuscript.

T.S. supervised the research and was responsible for this manuscript.

## Competing interests

The authorsdeclear no competing interests.

**Materials & Correspondence**

## Methods

### Data reporting

No statistical methods were used to predetermine sample size. All experiments were randomized, and the investigators were partially blinded to allocation during experiments and data analysis.

### Mice

All mice were bred in-house under standard conditions in the animal facility of Biological Resources Laboratory (BRL) at the University of Illinois at Chicago (UIC). All the colonies were SFB-positive in the BRL. C57BL/6J (Jax: 000664), *Rag2*^*-/-*^ (JAX: 008449), C57BL/6-Tg (Tcra,Tcrb) 2Litt/J as the SFB TCR^7B8^ Tg (JAX: 027230), C57BL/6-Tg (Tcra,Tcrb) 3Litt/J as the SFB TCR^1A2^ Tg (JAX: 028204), CD45.1 B6.SJL-Ptprca Pepcb/BoyJ (JAX: 002014) mice were originally purchased from The Jackson Laboratory before being bred in-house. All mouse experiments were approved by and performed in accordance with UIC’s Animal Care Committee (ACC) at UIC College of Medicine. Almost all phenotypes were compared using littermate controls and the resultant data were combined in most experiments as indicated. No mice were excluded from analyses unless clearly indicated. All *in vivo* transfer experiments were performed using 6-to 16-week-old animals (both males and females). SFB TCR^7B8^ *Rag2*^*-/-*^ mice, SFB TCR^1A2^ *Rag2*^*-/-*^, and their littermate controls were monitored until reaching 35 weeks of age.

### SFB mono-colonized mouse establishment

All germ-free (GF) and gnotobiotic mice were bred in-house in a satellite GF/gnotobiotic animal room in the Biomedical Research Building at the UIC. The SFB mono-colonization colony in the GF room is routinely tested by qPCR (every month) and by both anaerobic and aerobic culture testing (every week) to confirm a lack of any commensal microbes other than SFB. GF *Rag1*^*-/-*^ and *Rag2*^*-/-*^ mice were originally purchased from the University of Michigan and North Carolina GF facilities, respectively before being bred in-house under GF conditions. Original SFB-mono-colonized C57BL/6 mice were generated via oral gavage with the cecal contents of SFB mono-colonized mice provided by Yakult Inc. To prepare SFB mono-colonized *Rag1*^*-/-*^ and *Rag2*^*-/-*^ mice, GF *Rag1*^*-/-*^ and *Rag2*^*-/-*^ mice were transferred to an isolator housing SFB mono-colonized C57BL/6 for least 2-3 weeks prior to experimental use. After this transfer, the dirty bedding of SFB mono-colonized mice was added to the cages housing these *Rag1*^*-/-*^ and *Rag2*^*-/-*^ mice. This bedding addition was performed at least three times over two weeks. SFB colonization of these animals was confirmed by qPCR, immunological phenotyping, or microscopic observation using smashed fecal pellets.

### Naïve T cell transfer of SFB-specific TCR Tg cells into Rag-deficient hosts

Spleens and lymph nodes were collected from TCR^7B8^ or TCR^1A2^ Tg mice. Cells from these tissues were dissociated using 100 μm cell strainers, and red blood cells were lysed using ACK Lysing Buffer. Naïve CD4 T cells were presorted with the MojoSort™ Mouse CD4 Naïve T cell Isolation kit (BioLegend) based on provided directions. The magnetically sorted TCR^7B8^ or TCR^1A2^ naïve CD4 T cells were incubated with Fc Block (1:50; Tonbo 70-0161-M001: Purified Anti-Mouse CD16/CD32 (Fc Shield) (Clone:2.4G2)) and Goat Anti-Armenian Hamster IgG H+L (1:50; Jackson ImmunoResearch: 127-005-099) in RPMI 10% FCS for 15 minutes at 4°C, further stained with DAPI, anti-CD4-APC-Cy7 (BD Biosciences 552051: Clone: GK1.5), anti-CD8a-PE-Cy7 (Invitrogen 25-0081-82: Clone: 53-6.7, eBioscience), anti-CD62L-PE (Invitrogen 12-0621-82: Clone: MEL-14), anti-CD44-PerCP5.5 (Invitrogen 45-0441-82: Clone: IM7, eBioscience), anti-TCRv β14-FITC (BD Biosciences 553258: Clone: 14-2), and anti-CD25-APC (Invitrogen 17-0251-81: Clone: PC61.5, eBioscience), and then were purified as DAPI^-^, CD4^+^, CD8^-^, TCRβ+, CD62L^hi^, CD44^low^, CD25^-^, TCRVβ14+ CD4+ naïve T cells from spleen and lymph nodes of TCR Tg mice using the MoFlo Astrios or MoFlo Astrios EQ (Beckman) at the Flow Cytometry Core in the UIC Research Resources Center (RRC). After sorting, sorting efficiency was confirmed, and cells exhibiting > 99% purity for appropriate gating were used for the transfer experiments. Cells were counted with a hemocytometer, and 100,000 naïve T cells were administered to recipient mice via intravenous retro-orbital (RO) injection (100 μl total volume). Mice were euthanized via the intraperitoneal (i.p.) injection of ketamine and xylazine. Naïve T cells isolated from female donors were introduced into only female hosts. Naïve T cells isolated from male donors were introduced into male hosts in most cases, but were introduced into female donors in some cases. The injected hosts were monitored closely for 2 days, and their body weight and disease scores were checked at least every week until mice were used for tissue collection.

### Ampicillin treatment-mediated SFB removal

Based on our 3+ years of experience in the UIC BRL animal facility, all tested mice housed in our animal room are SFB-positive. To eliminate SFB from their commensal community, we treated these mice with 1 g/L Ampicillin in the drinking water for at least 2 weeks before the transfer of TCR^7B8^ naïve T cells into SFB-negative hosts. SFB colonization in Amp-treated hosts was tested via the collection of fecal pellets from random mice for microscopic observation and qPCR analyses using SFB-specific primer sets.

### Neurological phenotype scoring

Neurological phenotype scoring was performed as described in an article by Guyenet SJ. Et al.^67^ with minor modifications. Briefly, scoring consisted of 4 tests: A) Ledge test, B) Hindlimb clasping scores, C) Gait assay, and D) Kyphosis scoring. Scoring was performed weekly. The maximum possible score in each category was 3, with a maximum possible total neurological phenotype score of 12. Animals were euthanized if they exhibited scores of 10 or higher, or if their body weight fell to below 80% of baseline (day 0) values.

A. Ledge test: Lift the mouse from the cage and place it on the cage’s ledge. Score 0: A healthy mouse will typically walk along the ledge without losing its balance, and will lower itself back into the cage gracefully, using its paws. If the mouse loses its footing while walking along the ledge, but otherwise appears coordinated, it receives a score of 1. If it does not effectively use its hind legs, or lands on its head rather than its paws when descending into the cage, it receives a score of 2. If it falls off the ledge, or nearly so, while walking or attempting to lower itself, or shakes and refuses to move at all despite encouragement, it receives a score of 3.

B. Hindlimb clasping: Observe the mouse’s hindlimb position for 10 seconds. If the hindlimbs are consistently splayed outward, away from the abdomen, it is assigned a score of 0. If one hindlimb is retracted toward the abdomen for more than 50% of the time suspended, it receives a score of 1. If both hindlimbs are partially retracted toward the abdomen for more than 50% of the time suspended, it receives a score of 2. If its hindlimbs are entirely retracted and touching the abdomen for more than 50% of the time suspended, it receives a score of 3.

C. Gait: Remove the mouse from its cage and place it on a flat surface with its head facing away from the investigator. Observe the mouse from behind as it walks. If the mouse moves normally, with its body weight supported on all limbs, with its abdomen not touching the ground, and with both hindlimbs participating evenly, it receives a score of 0. If it shows a tremor or appears to limp while walking, it receives a score of 1. If it shows a severe tremor, severe limp, lowered pelvis, or its feet point away from the body during locomotion (“duck feet”), it receives a score of 2. If the mouse has difficulty moving forward and drags its abdomen along the ground, it receives a score of 3.

D. Kyphosis: Remove the mouse from its cage and place it on a flat surface. Observe it as it walks. If the mouse is able to easily straighten its spine as it walks and does not have persistent kyphosis, it receives a score of 0. If the mouse exhibits mild kyphosis but is able to straighten its spine, it receives a score of 1. If it is unable to straighten its spine completely and maintains persistent but mild kyphosis, it receives a score of 2. If the mouse maintains pronounced kyphosis as it walks or while it sits, it is assigned a score of 3.

Disease incidence was defined as 1) when a mouse first exhibits a score of 1 and maintains that score for 2+ weeks, or 2) when a mouse first exhibits a score of 2.

### Murine tissue collection for mononuclear cell isolation

To avoid potential peripheral blood contamination, mice were perfused with 50 ml of PBS containing 5 mM EDTA following euthanasia achieved via the i.p. administration of ketamine and xylazine. After perfusion, mLNs, spleen, small intestine, large intestine, liver, lung, brain, and spinal cord samples were collected in this order. The collected tissues were immediately stored in the ice-cold CM10 (RPMI 10% FBS,1x MEM, 10mM HEPES, 1mM Sodium Pyruvate, 1000 U/mL Penicillin, and 1000 mg/mL Streptomycin) and processed for the isolation of mononuclear cells described below.

### Pulmonary and hepatic mononuclear cell isolation

After perfusion, murine lung and liver tissue samples were transferred into 2 ml tubes containing 1 mL of digestion media (1 mg/mL Collagenase D (Sigma 11088882001: Collagenase D from Clostridium histolyticum), 100 ug/mL DNase (Sigma DN25: Deoxyribonuclease I from bovine pancreas), and 0.01 U/mL Dispase (Worthington Biological Corporation LS02104) in CM10). These tissues were minced with scissors in digestion media and transferred into 50 mL conical tubes. The remaining minced tissues in 2 ml tubes were collected by washing the tube with 6 mL of digestion media and all tissues were combined into one 50 mL conical tube. The tissues were then digested by vigorously shaking at 37°C (225-235 rpm) for 30 min. Tissues were then added to 14 mL of ice-cold CM10 to stop/delay reactions, after which they were passed through a 100 μM cell strainer and mononuclear lymphocytes were isolated using a Percoll (Cytiva 45001747) density gradient of 40%:80%. After the aspiration of the unnecessary cells and tissues in the upper 40% Percoll layer, lymphocytes were collected from the intermediate interface layer, washed with ice-cold CM10, and counted using Trypan Blue and a Countess II Automated Cell Counter (Fisher Scientific).

### Isolation of mononuclear cells from the brain and spinal cord

After perfusion, murine brain and spinal cord tissues were transferred into 2 mL tubes containing 1 mL of CNS digestion media (2.5 mg/mL Collagenase D and 100 μg/Ml DNase in CM10). These tissues were minced with scissors in digestion media and transferred into 50 mL conical tubes. The remaining minced tissues in 2 mL tubes were collected by washing the tube with 6 mL of digestion media and all tissues were combined into one 50 mL conical tube. The tissues were then digested by vigorously shaking at 37°C (225-235 rpm) for 30 min. Tissues were then physically digested by passing them through 21G needles 15-20 times, and the resultant tissue solutions were added 14 mL of ice-cold CM10 to stop/delay reactions. Tissues were then passed through a 40 μM cell strainer and mononuclear lymphocytes were isolated using a Percoll density gradient of 40%:80%. After the aspiration of the unnecessary cells and tissues in the upper 40% Percoll layer, lymphocytes were collected from the intermediate interface layer, washed with ice-cold CM10, passed through a 40 μM cell strainer, and counted using Trypan Blue and a Countess II Automated Cell Counter.

### Isolation of intestinal lamina propria (LP) mononuclear cells

LP mononuclear cells were isolated from the ileum (last 1/3 of the small intestine and large intestine including the cecum) following the procedure described in a previous study ^30^. Briefly, all the chains of mLNs were first collected or removed from the mice and the small intestine (SI) was separated from the large intestine (LI). The SI was further divided into three equal parts: duodenum (first 1/3 connecting to the stomach), jejunum (middle 1/3), and Ileum (last 1/3 connecting to the cecum). Attached fatty tissue was removed from the intestines. Tissues were cut into 3-4 pieces (5-8 cm), then placed in PBS containing 1 mM DTT (Sigma) and 5 mM EDTA and were incubated at 37°C on a shaker with vigorous shaking (225-235 rpm) for 20 min. Then, the incubated tissues were transferred into fresh PBS 5 mM EDTA and further shaken at 37°C on a shaker with vigorous shaking (225-235 rpm) for 10 min. Subsequently, tissues were washed in RPMI supplemented with 2% FBS, to remove DTT and EDTA. The incubated tissues were then minced by scissors in the 2ml tube and further digested in digestion media (1 mg/mL Collagenase D, 100 ug/mL DNase, and 0.01U/mL Dispase in CM10) at 37°C on a shaker (225-235 rpm) for 30 minutes for the ileum and 45 minutes for the LI. Tissues were then added to 14 mL of ice-cold CM10 to stop/delay reactions, after which they were passed through a 100 μM cell strainer and mononuclear lymphocytes were isolated using a Percoll density gradient of 40%:80%. After the aspiration of the unnecessary cells and tissues in the upper 40% Percoll layer, lymphocytes were collected from the intermediate interface layer, washed with ice-cold CM10, and counted using Trypan Blue and a Countess II Automated Cell Counter.

### Isolation of mononuclear cells from the spleen and mLNs

Spleen and mLNs were excised and stored on ice in CM10, after which they were dissociated by using the plunger from a 3mL syringe and pressing the tissues through a 70μM cell strainer, washing them with 5 mL of CM10. Spleen and mLN cells were then separated with a Percoll density gradient (40%:80%) to isolate mononuclear lymphocytes as above. After collection of the lymphocyte interface layer, the cells were washed with CM10 and counted using Trypan blue and a Countess II automated cell counter.

### Detection of T cell cytokine production

After the isolation of mononuclear cells from tissues, up to 1.0 million mononuclear cells per sample were plated in a 96-well round-bottom plates. These cells were restimulated for 4 hours at 37°C with CM10 containing 50 ng/mL PMA (Sigma), 500 ng/mL Ionomycin (Sigma), and 1x GolgiPlug (BD 555029). Following restimulation, the cells were washed with CM10 and incubated with Fc Block (1:50; Tonbo 70-0161-M001: Purified Anti-Mouse CD16/CD32 (Fc Shield) (Clone:2.4G2)) and Goat Anti-Armenian Hamster IgG H+L (1:50; Jackson ImmunoResearch: 127-005-099) for 15 minutes at 4°C. The cells were washed with CM10 and stained for extracellular markers of interest using antibodies including anti-CD3e-PerCP5.5 (Invitrogen 45-0031-82: Clone: 145-2C11 eBioscience), anti-TCRβ-APC ef780 (Invitrogen 47-5961-82: Clone: H57-597, eBioscience), anti-MHC II-BV510 (BD Biosciences 742893: Clone: M5/114.15.2 BD Optibuild), anti-CD4-BV605 (BD Biosciences 563151: Clone: RM4-5), and anti-CD8a Super Bright 702 (Invitrogen 67-0081-82: Clone: 53-6.7, eBioscience) for 30 minutes in the dark at 4°C (all antibodies were diluted 1:200 with CM10). The cells were then washed with PBS twice and stained with Aqua Live/Dead stain (Invitrogen L34966) (1:200) for 30 minutes in the dark at 4°C. The cells were washed with PBS and fixed with BD Cytofix/Cytoperm Fixation/Permeabilization Solution Kit (BD 555028) Kit for 30 minutes in the dark at 4°C. The cells were then permeabilized using BD fixation Kit Permeabilization Buffer and sometimes stored at 4°C over night. Finally, cells were stained for intracellular markers using antibodies including anti-IFNγ-AF488 (Invitrogen 53-7311-82: Clone: XMG1.2, eBioscience), anti-IL-17A-AF700 (BioLegend 506914: Clone: TC11-18H10.1), anti-IL-22-APC (Invirtogen 17-7222-80: Clone: IL22JOP, eBioscience), anti-TNFa-BV421 (BD Bioscience 563387: Clon MP6-XT22) or anti-TNFa-PE (Invirtogen 14-7321-81: Clon MP6-XT22, eBioscience), anti-GM-CSF-PE (Invitrogen 12-7331-82: Clone: MP1-22E9, eBioscience), and anti-IL-13-PECy7(Invitrogen 25-7133-82: Clone: eBio13A, eBioscience) (all 1:200) for 30 minutes in the dark at 4°C. Cells were washed using BD permeabilization buffer and filtered through 40 or 70 μM cell strainers before running them on an Attune NxT Cytometer (Invitrogen).

### Detection of cell surface markers and intracellular proteins in T cells

After isolation of mononuclear cells from tissues, 1.0 million mononuclear cells per sample was plated in 96-well round-bottom plates. These cells were immediately blocked using Fc Block (1:50; Tonbo 70-0161-M001: Purified Anti-Mouse CD16/CD32 (Fc Shield) (Clone:2.4G2)) and Goat Anti-Armenian Hamster IgG H+L (1:50; Jackson ImmunoResearch: 127-005-099) for 15 minutes at 4°C. They were then washed with CM10 and extracellular staining was performed using anti-CD44-PerCP Cy5.5 (Invitrogen 45-0441-82: Clone: IM7, eBioscience), anti-TCRβ-APC ef780 (Invitrogen 47-5961-82: Clone: H57-597, eBioscience), anti-MHC II-BV510 (BD Biosciences 742893: Clone: M5/114.15.2 BD Optibuild), anti-CD4-BV605 (BD Biosciences 563151: Clone: RM4-5), and anti-CD8a Super Bright 702 (Invitrogen 67-0081-82: Clone: 53-6.7, eBioscience) for 30 minutes in the dark at 4°C. Cells were washed using PBS and stained with Aqua Live/Dead stain (Invitrogen L34966) for 30 minutes in the dark at 4°C (all antibodies were diluted 1:200 with CM10). The cells were then washed with PBS and fixed using an eBioscience Fixation Kit (Invitrogen 00512343 and 00522356) for 30 minutes in the dark at room temperature. The cells were permeabilized with eBioscience Permeabilization Buffer (Invitrogen 00833356) and stained using antibodies specific for intracellular markers including anti-Ki67-FITC (Invitrogen 14-5698-82: Clone: SolA15, eBioscience) (1:200), anti-FoxP3-APC (Invitrogen 17-5773-82: Clone: FJK-16s, eBioscience) (1:200), anti-RORγt BV421 (BD Biosciences 562894: Clone: Q31-378) (1:150), anti-GATA3-PEcy7 (Invitrogen 25-9966-42: Clone: TWAJ, eBioscience) (1:50), and anti-Tbet-PE (Invitrogen 12-5825-82: Clone: 4B10, eBioscience) (1:25) for 30 minutes in the dark at room temperature. The cells were again washed with eBioscience permeabilization buffer and passed through a 40-70 μM cell strainer before running on Attne NxT Cytometer (Invitrogen).

### ELISA-mediated detection of lipocalin 2 (LCN2) in fecal pellets^68^

Fresh fecal pellets were collected directly from mice and stored at -80°C until all the samples for that experimental group were gathered. The frozen fecal pellets were reconstituted in PBS containing 0.1% Tween 20 (100 mg of pellets in 1 ml of PBS0.1% Tween20). The fecal pellets were mixed for 20 min on shaker to produce a homogenous fecal suspension. These suspensions were then centrifuged for 10 min at 12,000 rpm at 4°C. Clear supernatants were carefully collected and stored at -20°C until analysis. Fecal LCN2 levels in the supernatant fraction were estimated using a Duoset murine LCN-2 ELISA kit (R&D Systems DY1857-05, Minneapolis, MN, USA) based on provided directions.

### In vitro TCR^7B8^ and TCR^1A2^ stimulation using purified and crude peptides

In vitro naïve T cell stimulation assay in an article by Leonard JD et al.^69^ was modified and optimized for TCR^7B8^ naïve T cells. Both crude (∼50% purity) and purified (>95% purity; TFA removed) peptides were synthesized by GenScript USA, Inc. (NJ USA), reconstituted with ultrapure water or DMSO in the laboratory, and stored at -20°C for several months or -80°C for long-term storage. Bone marrow was collected from CD45.1/CD45.1 C57BL6 mice, red blood cells were removed using ACK lysis buffer (Lonza), and cells were then cultured in 10 cm coated dishes for 7-8 days in complete cell culture media (RPMI 10% FBS, 1x MEM, 10 mM HEPES, 1 mM Sodium Pyruvate, 1000 U/mL Penicillin, 1000 mg/mL Streptomycin, and 1x β-ME). For selective differentiation into the dendritic cells, 15 ng/ml of recombinant murine GM-CSF (PEPROTECH: 315-03) was added on day 0, day 3, and day 7 for bone marrow cell culture. On day 8-9 of culture, 2-4×10^4^ bone marrow-derived DCs (BMDCs) were re-plated in 96-well round-bottom plates and stimulated with 1.0 μg/ml LPS (Sigma L2762: Escherichia coli 26:B6) and different concentrations of peptides for 24 hours. For *in vitro* stimulation assays, naïve CD4 T cells were magnetically isolated from the spleen and LNs of *Rag2*^*-/-*^ TCR^7B8^ Tg or TCR^1A2^ Tg mice on *CD45*.*2/CD45*.*2* background using the MojoSort™ Mouse CD4 Naïve T cell Isolation kit (BioLegend). Then, same number of magnetically sorted naive T cells were co-cultured with equal numbers of BMDCs supplemented with 20 ng/ml of recombinant mouse IL-2 (PEPROTECH: 212-12) for 24-72 hours. In some cases, to block MHC class II, 10 μg/ml of unconjugated Anti-MHCII (BD Bioscience 556999: I-A/I-E Rat anti-Mouse, Unlabeled Clone: M5/114.15.2.) were added 1 hour before naïve T cell addition to the co-culture system. Naïve T cell activation was examined by the expression of CD25 (anti-CD25-APC (Invitrogen)) and CD69 (anti-CD69-AF700 (BD biosciences 561238: Clone: H1.2F3)) on DAPI^-^, TCRβ+ (anti-TCRβ-APC ef780 (Invitrogen 47-5961-82: Clone: H57-597, eBioscience)), CD4+ (anti-CD4-FITC (Tonbo 350042U500: Clone: RM4-5)), CD45.2+ (anti-CD45.2-PE-Cy7 (Tonbo 600454U100: Clone: 104)) TCR Tg T cells via FACS (all was diluted 1:200). In some experiments, magnetically sorted naïve T cells were stained using CFSE (Thermo C34554: CellTrace™ CFSE Cell Proliferation Kit for flow cytometry)) just before the co-culture with BMDCs, and CFSE dilution, Ki67 expression (anti-Ki67-eFluor 450 (Invitrogen 48-5698-82: Clone:SolA15)), Nur77 expression (anti-Nur77-PE (Invitrogen 12-5965-82: Clone: 12.14, eBioscience)), CD25 (anti-CD25-APC (Invitrogen)), and CD69 (anti-CD69-AF700 (BD biosciences 561238: Clone: H1.2F3)) were assessed in Aqua^-^, TCR+, CD4+, CD45.1+ TCR Tg T cells via FACS to measure proliferation. Ki67 and Nur77 were stained following the extracellular staining, Aqua (Thermo L34966) staining, and fixation step using an eBioscience Fixation Kit (Invitrogen 00512343 and 00522356) for 30 minutes in the dark at room temperature. The cells were washed with eBioscience permeabilization buffer and passed through a 40-70 μM cell strainer before running on Attne NxT Cytometer (Invitrogen).

### Brain immunohistochemical (IHC) staining

Mice were initially anesthetized using the isoflurane drop method and were subsequently kept under deep anesthesia via continuous exposure to isoflurane. Mice were then transcardially perfused with 30-40 mL of ice-cold PBS, followed by 20 mL of ice-cold 4% PFA in PBS. After perfusion, the brain was resected and placed in 4% PFA overnight at 4°C for post-fixation before being transferred to 30% sucrose in PBS for 48 hours. Processed brains were then halved at the sagittal midline cut into 50 μm sagittal sections in ice-cold PBS using a vibratome (Vibratome Series 1000, TPI). Sections were stored at -20°C in cryoprotectant solution (30% glycerol, 30% ethylene glycol in PBS) until immunostaining was performed.

All sample immunostaining for Iba1 (catalog no.: 019-19741; AB_839504) was conducted in parallel to avoid interexperimental variation. Briefly, 7-8 representative sections from each brain were initially washed in TBS for 10 min prior to transfer into immunostaining baskets and incubation in Tris-EDTA antigen retrieval solution (10 mM Tris, 1 mM EDTA, pH 9.0) at 90°C for 30 minutes. Sections were then allowed to equilibrate to room temperature in the same antigen retrieval solution before being washed once in TBS containing 0.1% Triton X-100 (TBSX) and permeabilized for 30 minutes in 0.5% TBSX at room temperature with shaking. Next, sections were blocked in 5% normal goat serum plus 2% BSA in TBSX for 2 hours at room temperature. Sections were then incubated with primary anti-Iba-1 (1:1000 in antibody solution; 2% BSA in TBSX) for 48 hours at room temperature with shaking before being washed 4 times, 10 minutes each in TBSX Sections were next incubated with secondary antibody (1:500 goat anti-rabbit Alexa Fluor 488; catalog no.: A32731; AB_2633280) overnight protected from light. Sections were then washed as above before being mounted onto slides with VectaShield containing DAPI (Vecta Laboratories H1200-10: VECTASHIELD Antifade Mounting Medium with DAPI). Slides were imaged and data were acquired using a confocal microscope (LSM710, Zeiss) or a Keyence inverted fluorescence microscope (BZ-X810, Keyence) under identical imaging conditions and settings.

### Spinal cord IHC staining

After the mice were anesthetized using Ketamine and Xylazine, they were perfused with 40 mL of ice-cold PBS(-) and then with 30 mL of ice-cold 4% PFA (Electron Microscopy Science 15714) in PBS. Following perfusion and fixation, the spinal cord was placed in 4% PFA PBS for 30 minutes. Samples were then washed thrice with cold PBS(-) for 30 minutes. Then the tissues were further incubated in 30% sucrose in PBS overnight until the samples no longer floated to the top of the tube. Tissues were then removed from this sucrose solution and the spinal cord was cut into four equal parts. Tissues were placed in a cassette and embedded with O.C.T (Fisher Scientific 23-730-571: Scigen Tissue-Plus™ O.C.T Compound)) and frozen and stored at -80°C. When ready for staining, tissues are removed from storage, cut into 12-14 μm sections with a Cryostat, and placed on microscopy slides. Sections stored at -80°C prior to staining.

### Myelin staining of spinal cords with FluorMyelin^70^

Briefly, frozen tissue sections described in the section of Spinal cord IHC staining were permialized in PBT (PBS + 0.2% Triton X-100) for at least 20 min and then stained with FluoroMyelin™ Green fluorescent myelin stain solution or BrainStain™ Imaging Kit (Molecular Probes, Invitrogen, Eugene, OR) diluted 1:300 in PBS for 20 min. Slides were washed three times for 10 min each with PBT and mounted with an aqueous antifade mounting medium such as ProLong or ProLong Gold antifade reagent. Images were visualized by Keyence BZ-X710 microscope.

### IHC Staining of brain and spinal

Sectioned slides were removed from storage at -80°C and air-dried for 30 minutes at room temperature (RT). Following drying, the sections were incubated in 1x PBS for 30 minutes at RT to remove O.C.T. Slides are then blocked for 1 hour in a humidified chamber with 1 mL of PBS(-) containing 0.1% Tween20 and 10% bovine serum albumin (BSA) (GeminiBio 700-100P). After blocking, media was removed and replaced with 800 μL of PBS(-) containing 0.1% TritonX-100 (SIGMA T8787-250mL) and 1% BSA (GeminiBio 700-100P) with appropriately diluted primary antibodies, and slides were placed in a humidified chamber at 4°C overnight. The next day, slides were washed three times for 10 minutes each in PBS(-) containing 0.1% Triton X-100. Samples were then incubated with 800 μL of PBS (-) containing 0.1% Triton X-100 and diluted secondary antibodies for 1-2 hours at RT protected from light. Sections were then washed three times for 10 minutes each with PBS(-) containing 0.1% Triton X-100 in the dark. Two more washes were performed with PBS(-) in the dark. After washing was complete, mounting media (Vecta Laboratories H1200-10: VECTASHIELD Antifade Mounting Medium with DAPI) was applied and a coverslip was attached over the tissue. Slides were then imaged using a Keyence BZ-X710 microscope.

### Checkpoint inhibitor treatment

Seven days following naïve TCR^7B8^ CD4 T cell transfer into the *Rag2*^*-/-*^ hosts, 300 μg of anti-CTLA4 (BioXCell BE0164: Clone:9D9) and anti-PD-1 (BioXCell BE0146: Clone:RMP1-14) were injected into these animals (i.p.). Antibody injections were performed weekly until 7 weeks following naïve T cell transfer. Control animals were instead injected with 600 μg of control antibodies (BioXCell BE0086: mouse IgG2 isotype control, unknown soecificity) using the same injection schedule. At 4 weeks and 6 weeks following the transfer, fresh fecal pellets were collected and used for LCN2 detection. At 7-8 weeks following transfer, brain, spinal cord, spleen, ileum, and colon samples were collected for the analysis of cytokines derived from TCR^7B8^ CD4 T cells.

### Active EAE induction

We induced EAE using 8-to 12-week-old female mice that were subcutaneously immunization with a 100 μL emulsion of 100 μg of MOG35–55 peptide (MEVGWYRSPFSRVVHLYRNGK) (Thermo Fisher Scientific) in PBS with CFA containing 200 μg of Mycobacterium tuberculosis H37Ra (Difco). The peptide-injected mice further received intravenous injections of 400 ng of *B. pertussis* toxin (List Biological Laboratories) 0- and 2-days post-immunization. Mice were examined for clinical signs of EAE using the following scale, with 0.5-point gradations for intermediate presentation: 0, no signs; 1, flaccid tail; 2, hindlimb paresis; 3, hindlimb paralysis; 4, hindlimb and forelimb paralysis; and 5, moribund^71^.

### Statistical analyses

Statistical analyses were performed using the GraphPad Prism software suite (v.9.2.0) and Microsoft Excel. Statistical differences between two groups of data were evaluated using two-tailed unpaired Student’s t-tests (for normally distributed variables) or the two-tailed unpaired Mann-Whitney test (for non-normally distributed variables). For disease incidence, the two-tailed log-rank (Mantel–Cox) test was performed. P < 0.05 was the threshold of statistical significance. *P < 0.05, **P < 0.01, ***P < 0.001, #P < 0.0001, and N.S. Not Significant. All the statistical details for individual experiments are provided in the figure legends.

## Supplemental Movie 1

**Neurological phenotype in non-transferred *Rag2***^***-/-***^ **mouse**

Non-transferred SFB-colonized *Rag2*^*-/-*^ host 8 weeks following the transfer experiment co-housed with TCR7B8 Transferred SFB-colonized *Rag2*^*-/-*^ host. Neurological Score 0.

**Supplemental Movie 2**

**Neurological phenotype in TCR**^**7B8**^ **transferred *Rag2***^***-/-***^ **host with mild clasping**

1 × 10^5^ SFB TCR^7B8^ CD4 naïve T cells were retro-orbitally transferred into SFB-colonized *Rag2*^*-/-*^ hosts. Non-and TCR^7B8^-transferred *Rag2*^*-/-*^ hosts were co-housed after their weaning. Neurological Score at 8 weeks following the transfer: 3.

**Supplemental Movie 3**

**Neurological phenotype in TCR**^**7B8**^ **transferred *Rag2***^***-/-***^ **host with severe clasping**

1 × 10^5^ SFB TCR^7B8^ CD4 naïve T cells were retro-orbitally transferred into SFB-colonized *Rag2*^*-/-*^ hosts. Non-and TCR^7B8^-transferred *Rag2*^*-/-*^ hosts were co-housed after their weaning. Neurological Score at 8 weeks following the transfer: 6.

**Supplemental Movie 4**

**Neurological phenotype in TCR**^**7B8**^ **transferred Rag2**^***-/-***^ **host with severe clasping in both fore and hind limbs**

1 × 10^5^ SFB TCR^7B8^ CD4 naïve T cells were retro-orbitally transferred into SFB-colonized *Rag2*^*-/-*^ hosts. Non-and TCR^7B8^-transferred *Rag2*^*-/-*^ hosts were co-housed after their weaning. Neurological Score at 8 weeks following the transfer: 8.

