## Extended figures for "Commensal-specific CD4 T cells promote inflammation in the central nervous system via molecular mimicry"

Extended Data Figure 1

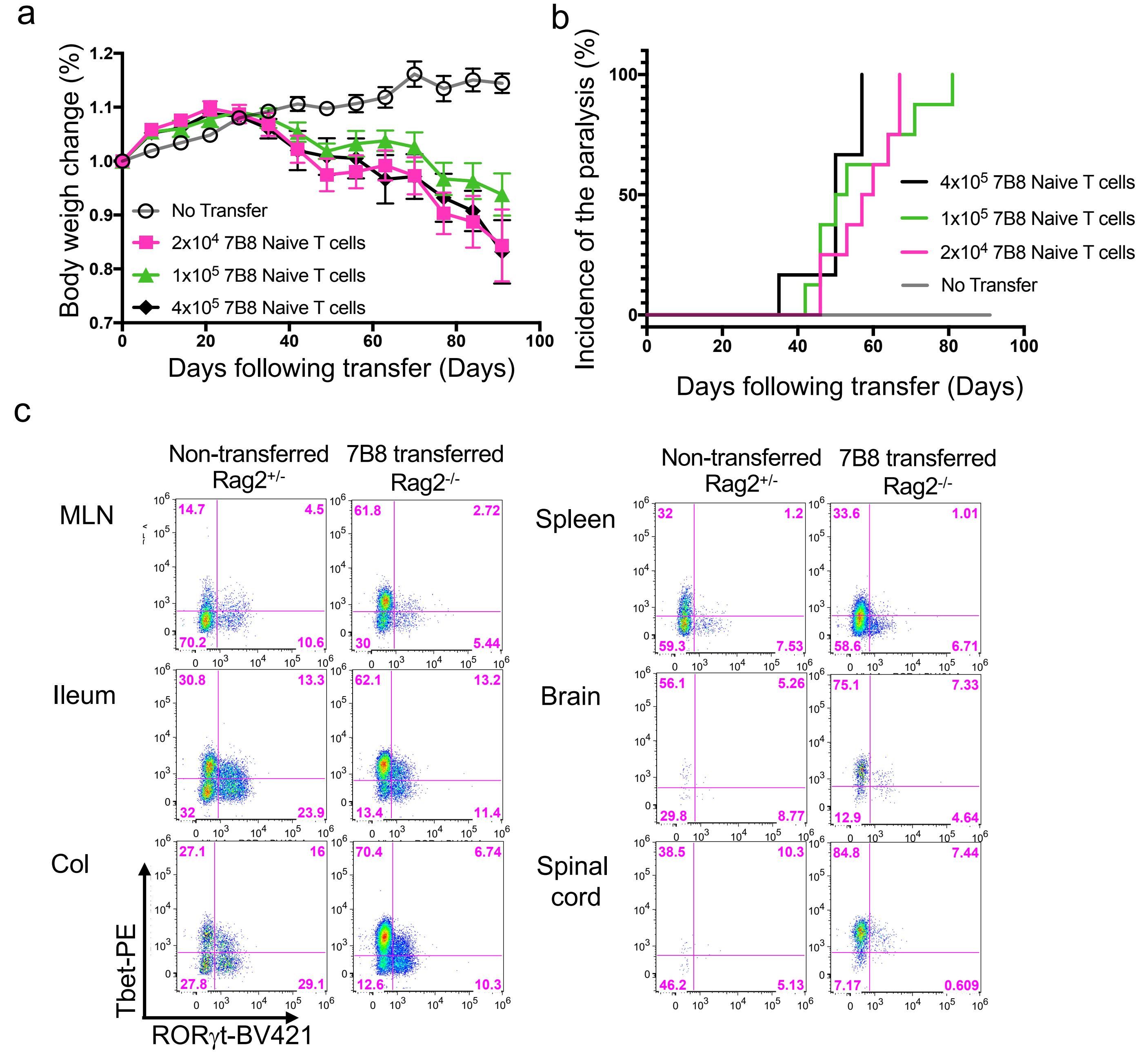


**Extended Data Fig. 1. Transferred SFB TCR^7B8 CD4 T cells expand and exhibit pathogenic activity in the intestines and CNS. a-c**, 2 x $10^4$, 1 x $10^5$, or 4 x $10^5$ SFB TCR^7B8 CD4 naïve T cells were retro-orbitally transferred into SFB-colonized *Rag2^-/-* hosts. After transfer, changes in body weight (**a**) and hind limb clasping incidence (**b**) were assessed. A one-time experiment with indicated numbers of replicates comprised primarily of littermates was conducted, n=8 (2 x $10^4$ transferred *Rag2^-/-* hosts), n=8 (1 x $10^5$ transferred *Rag2^-/-* hosts), n=6 (4 x $10^5$ transferred *Rag2^-/-* hosts), and n=8 (non-transferred *Rag2^-/-* hosts). Representative FACS plots of Tbet- and/or RORγt-expressing TCRβ⁺ CD4⁺ T cells in the mLN, ileum, colon, spleen, brain, and spinal cord of 1 x $10^5$ TCR^7B8 naïve T cells transferred *Rag2^-/-* hosts (n=3) at 8 weeks post-transfer (**c**). Non-transferred *Rag2^+/-* aged- and gender-matched controls were also used (n=3).

### Extended Data Figure 2

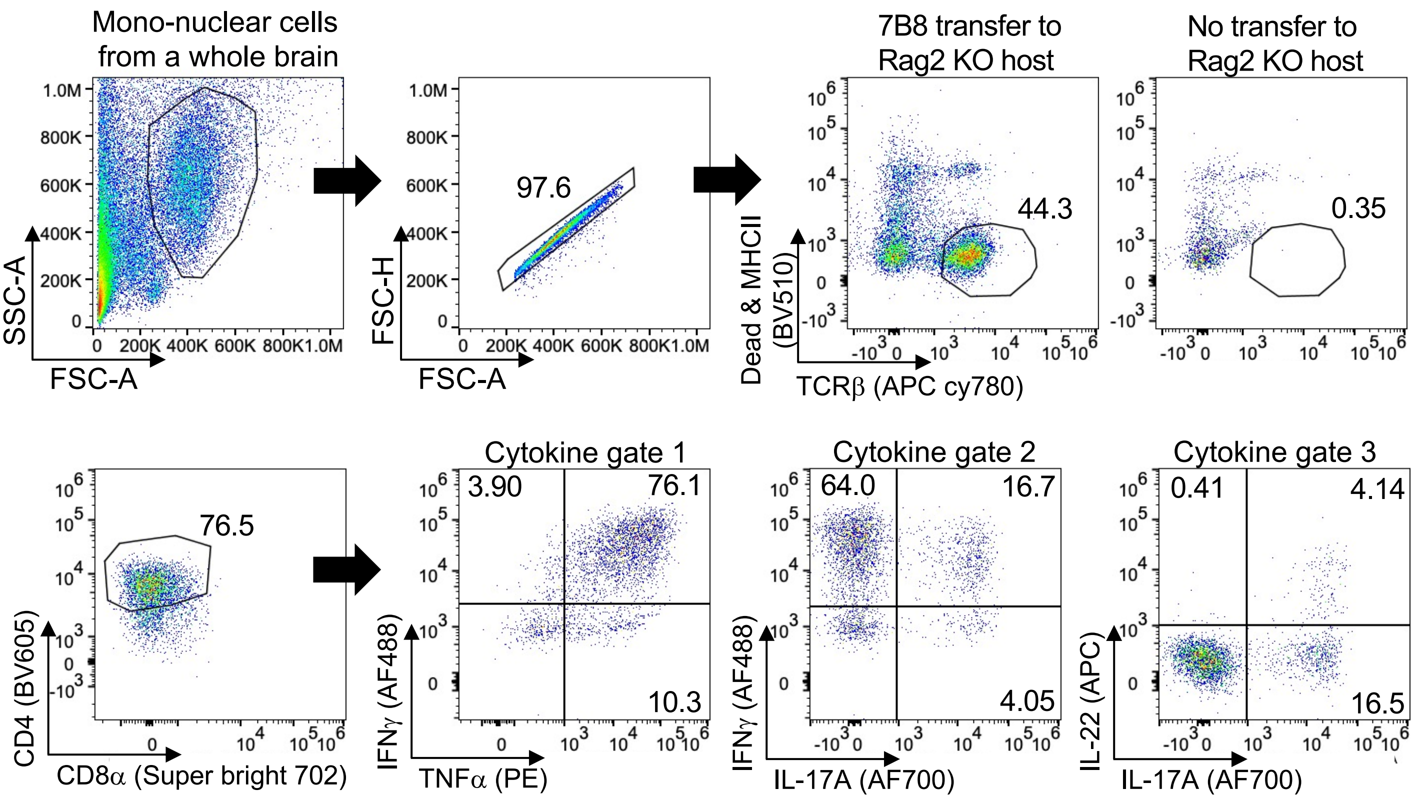


**Extended Data Fig. 2. Gating strategy for the analysis of TCR$^{7B8}$ CD4 T cells in tissues.** The gating strategy used to analyze the cytokine-producing T cells is shown. Following the transfer of 1 x $10^5$ SFB TCR$^{7B8}$ naïve CD4 T cells, lymphocytes were enzymatically collected from tissues and restimulated with PMA/Ionomycin and GolgiPlug for 4 hours. After the gating on FSC and SSC and eliminating doublet events, Dead cells and MHCII-expressing cells were excluded, and TCRβ$^+$, CD4$^+$, and CD8$^-$ T cells were further analyzed for the expression of the indicated cytokines (gate 1: IFNγ and TNFα, gate 2: IFNγ and IL-17A, and gate 3: IL-22 and IL-17A).

Extended Data Figure 3

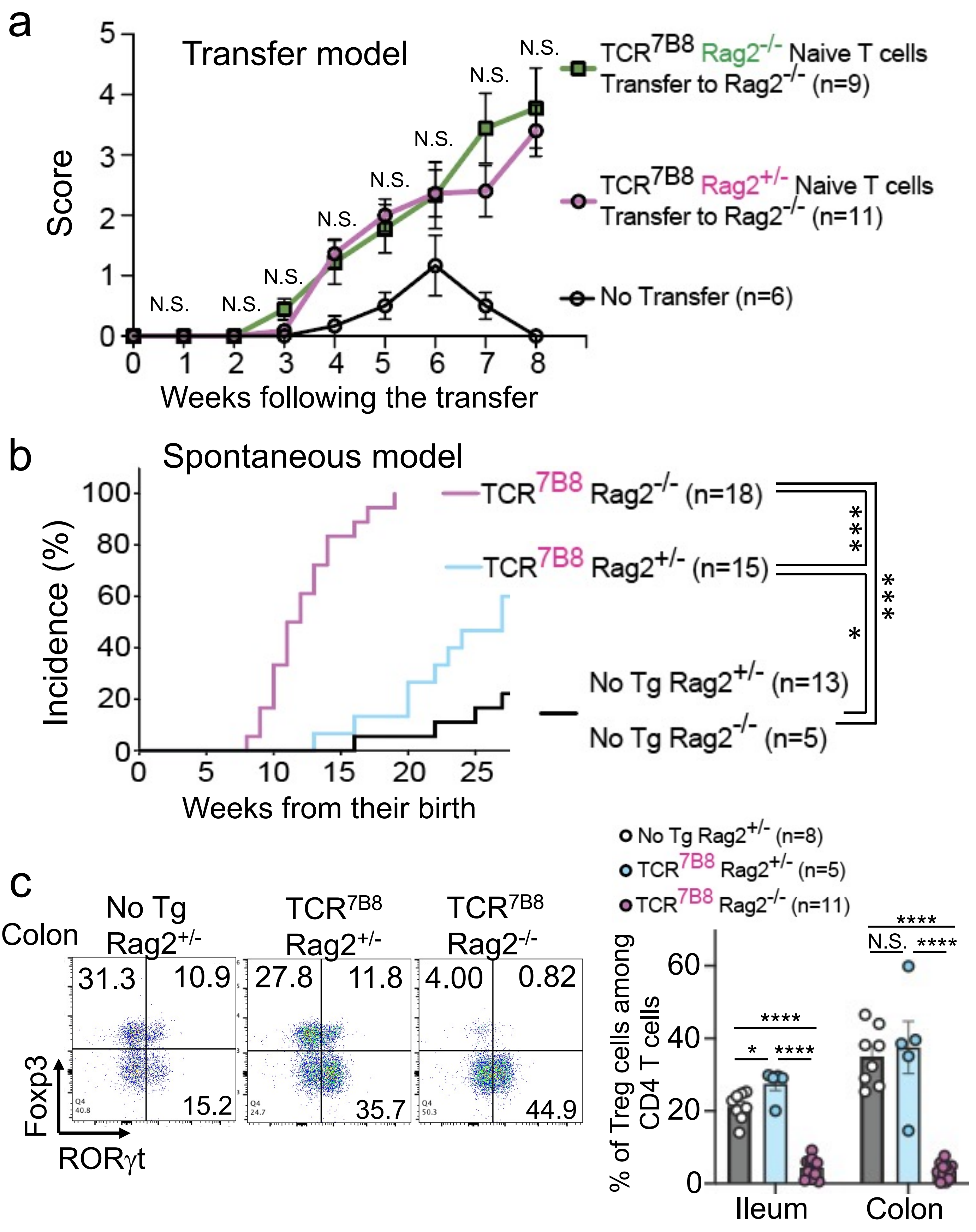

**Extended Data Fig. 3. Comparison of neurological phenotypes associated with TCR$^{7B8}$ on *Rag2*-sufficient and -deficient backgrounds. a,** SFB-colonized *Rag2*$^{-/-}$ hosts received a retro-orbital transfer of 1 x $10^5$ CD4 naïve T cells collected from SFB TCR$^{7B8}$ *Rag2*$^{+/-}$ or $^{-/-}$ donors. Neurological scores were monitored every week following the transfer. Two experiments were conducted and the scores from the indicated numbers of mice were pooled and reported as the mean ± SEM. Dots correspond to individual mice. **b**, The incidence of spontaneously developing neurological phenotypes for TCR$^{7B8}$ *Rag2*$^{-/-}$, TCR$^{7B8}$ *Rag2*$^{+/-}$, and No TCR$^{Tg}$ *Rag2*$^{-/-}$ and $^{+/-}$ mice. Littermates were compared and all the results were combined. Dots correspond to individual mice. **c**, Flow plots corresponding to Foxp3- and/or RORγt-expressing regulatory T cells (left) and the frequency of regulatory T cells (CD4$^+$, TCRβ$^+$, Foxp3$^+$, and RORγt$^-$ (peripheral Treg) plus CD4$^+$, TCRβ$^+$, Foxp3$^+$, and RORγt$^+$ (inducible Treg)) cells (right), in the indicated tissues from 30-35-week-old SFB TCR$^{7B8}$ *Rag2*$^{-/-}$, SFB TCR$^{7B8}$ *Rag2*$^{+/-}$, and non-TCR$^{Tg}$ *Rag2*$^{+/-}$ mice. Multiple experiments were conducted and the data from the indicated numbers of mice were pooled and reported as the mean ± SEM. Dots correspond to individual mice. Student's t-test. ****$P<0.0001$, ***$P<0.001$, **$P<0.01$, *$P<0.05$, N.S.; not significant ($P>0.05$).

### Extended Data Figure 4

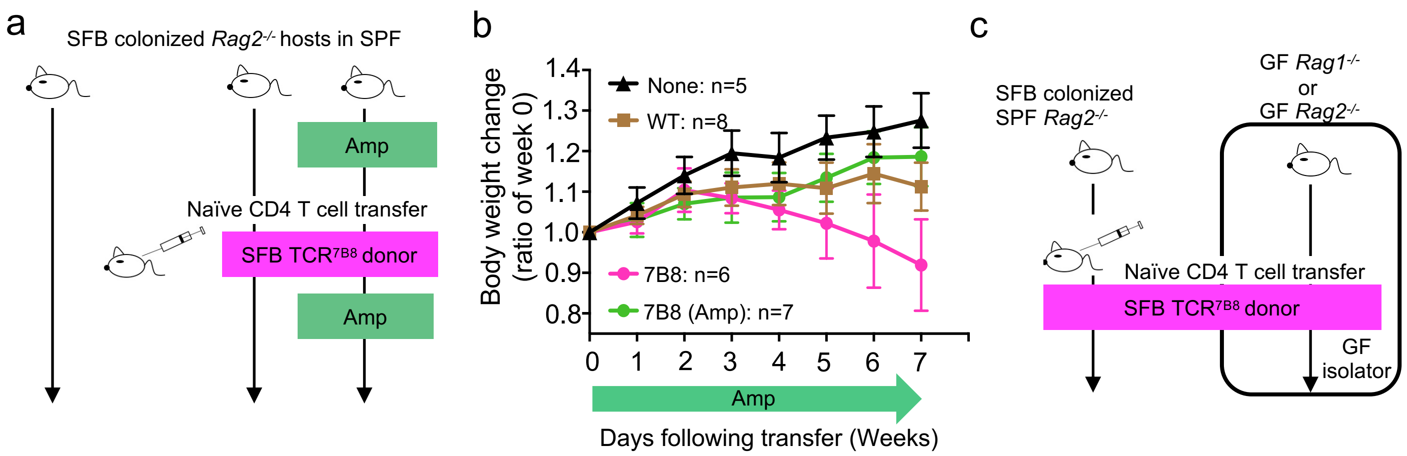


**Extended Data Fig. 4. Colonization of SFB is necessary and sufficient for TCR$^{7B8}$ naïve T cell-induced CNS inflammation. a**, Schematic representation of Fig. 1n-q. SFB-colonized *Rag2$^{-/-}$* hosts housed in the SPF facility were treated with 1 g/L Ampicillin in the drinking water for 7 days prior to the transfer of 1 x $10^5$ naïve TCR$^{7B8}$ CD4 T cells via RO injection. The ampicillin-containing water was changed every week for 8 weeks following the transfer. **b**, After transfer, changes in body weight were monitored every week. **c**, Schematic representation of Fig. 1n-q for SFB mono-colonization. Inside the GF isolator, 1 x $10^5$ TCR$^{7B8}$ naïve CD4 T cells were transferred (RO) into SFB mono-colonized Rag1 or *Rag2$^{-/-}$* hosts. Transferred hosts were kept inside the GF isolator for 8 weeks. These experiments were done twice, and all results were combined. The experiments for ampicillin treatment and SFB mono-colonization were conducted at the same time using the same donor cells.

Extended Data Figure 5

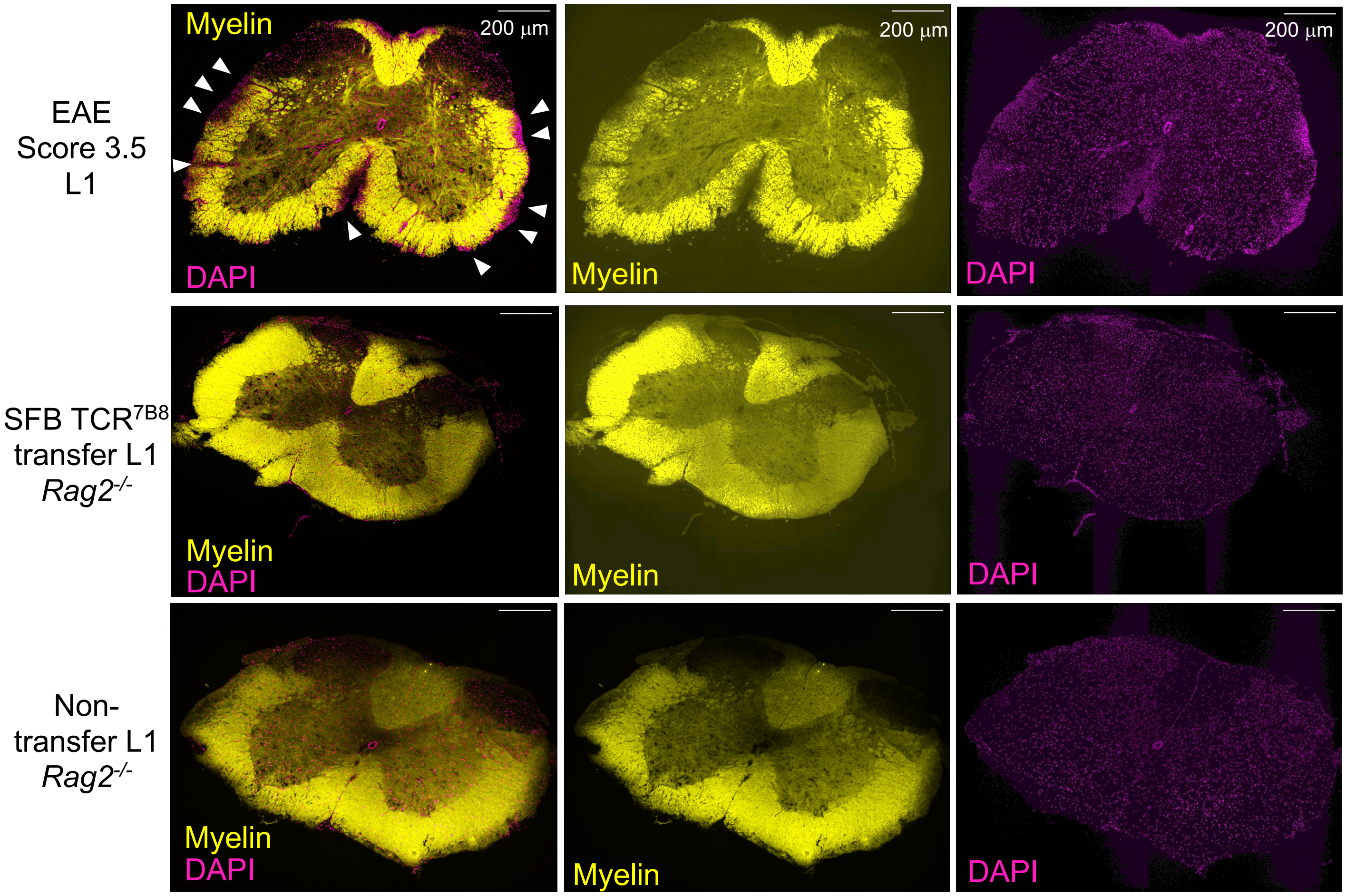


**Extended Data Fig. 5. Myelin disruptions were observed in EAE-induced hosts, but not in naïve TCR$^{7B8}$ CD4 T cell-transferred hosts.** Staining with FluorMyelin™ Green in the spinal cord (L1) of EAE-induced C57BL/6 WT (Top) and naïve TCR$^{7B8}$ CD4 T cell-transferred C57BL/6 *Rag2*$^{-/-}$ hosts (Middle), and non-transferred C57BL/6 *Rag2*$^{-/-}$ hosts (bottom). Stained myelin and cells (DAPI) were represented in yellow and magenta, respectively. Similar results were observed in 3 biological replicates (transferred and non-transferred *Rag2*$^{-/-}$ hosts). EAE scores were as indicated.

### Extended Data Figure 6

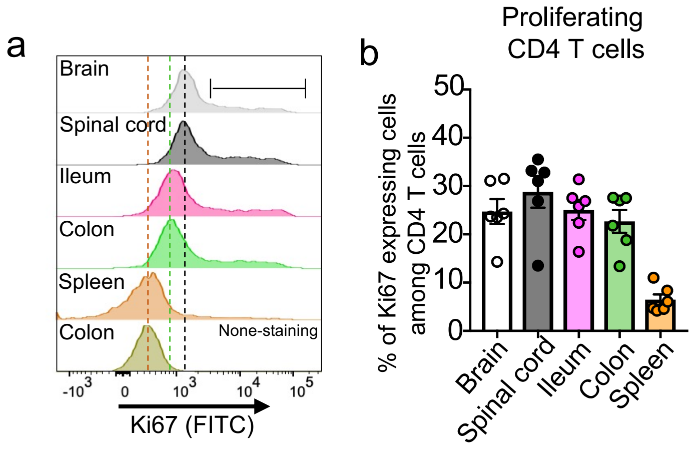


**Extended Data Fig. 6. Transferred SFB TCR$^{7B8}$ CD4 T cells were actively proliferating in the intestine and CNS of SFB TCR$^{7B8}$ *Rag2$^{-/-}$* mice. a,** Flow plots corresponding to Ki67-expressing TCRβ$^{+}$ CD4$^{+}$ CD8$^{-}$ T cells in the indicated tissues collected from 30-35-week-old SFB TCR$^{7B8}$ *Rag2$^{-/-}$* mice. **b**, Frequencies of Ki67-positive TCRβ$^{+}$ CD4$^{+}$ CD8$^{-}$ T cells in the indicated tissues. Three repeated experiments were performed with similar results. Littermates were used. Multiple experiments were conducted and the data from the indicated numbers of mice were pooled and reported as the mean ± SEM.

### Extended Data Figure 7

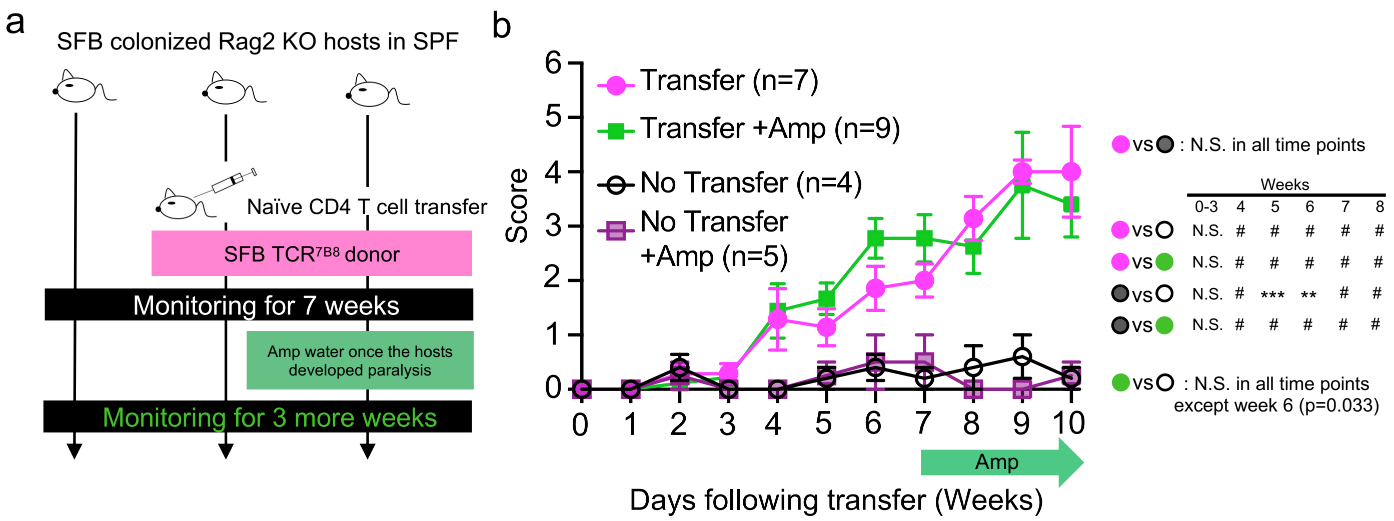


**Extended Data Fig. 7. Elucidation of therapeutic potential of CNS inflammation by Amp-treatment in hindlimb clasping transferred hosts. a**, Schematic overview of the approach to removing SFB after recipients had developed neurological phenotypes as a treatment intervention. SFB-colonized *Rag2*$^{-/-}$ hosts into which 1 x $10^5$ TCR$^{7B8}$ naïve CD4 T cells had been transferred were for neurological scores every week for 7 weeks. After all recipients had been confirmed to develop neurological phenotypes, 9 of the recipients were treated with ampicillin in the drinking water to eliminate SFB from the gastrointestinal tract. As controls, 7 of the recipient mice were instead given regular water. Non-transferred littermate controls were also monitored with or without ampicillin treatment. **b**, Hind limb clasping scores are shown together with the indicated numbers of mice used. Two independent experiments were conducted and the data from the indicated numbers of mice were pooled and reported as the mean ± SEM. Student's t-test. #P<0.0001, ***P<0.001, **P<0.01, N.S.; not significant (P>0.05).

#### Extended Data Figure 8

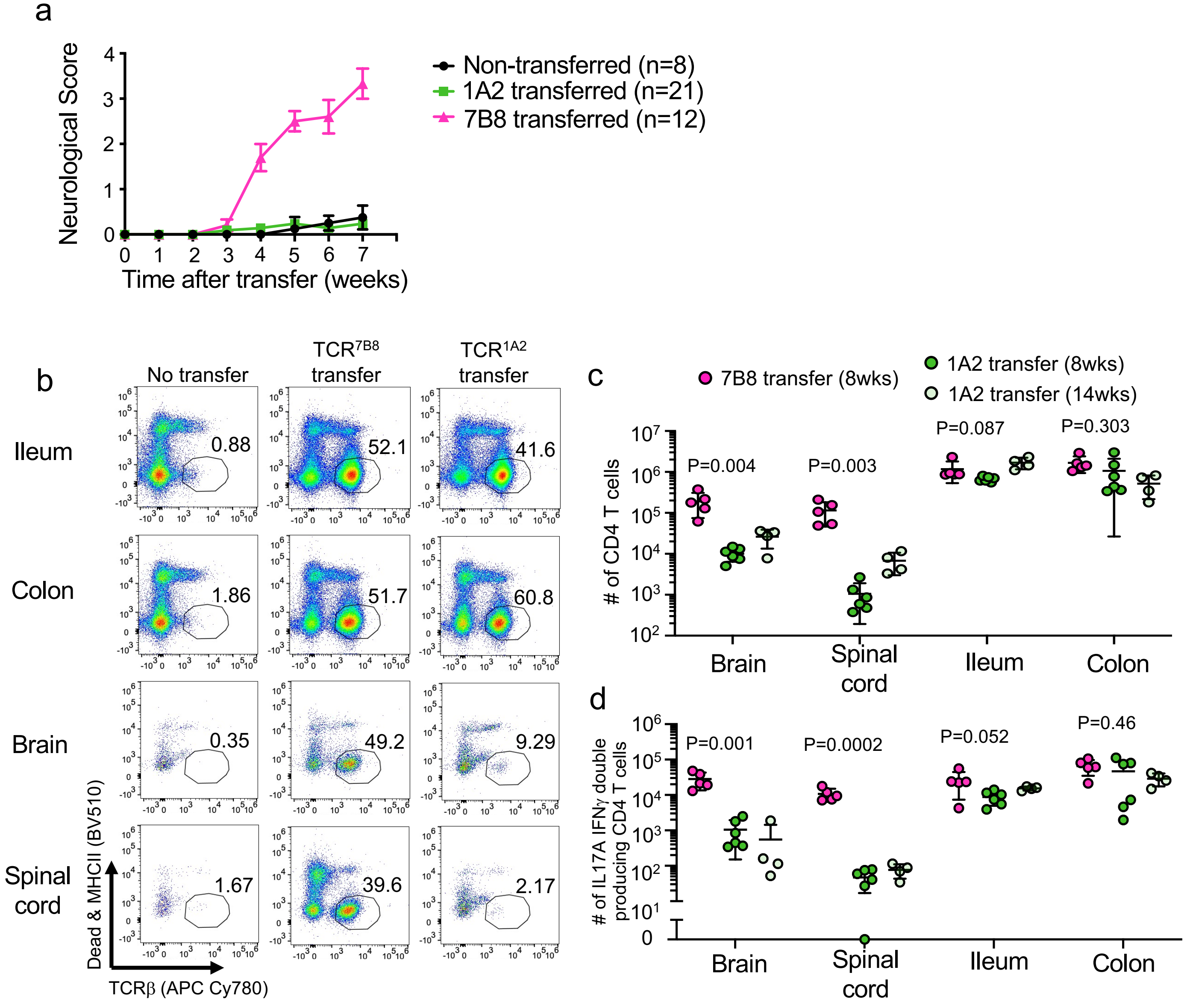


**Extended Data Fig. 8. The transfer of TCR7B8 naïve T cells but not TCR1A2 naïve T cells into *Rag2*$^{-/-}$ mice results in the development of CNS inflammation. a,** Neurological phenotype scores in *Rag2*$^{-/-}$ hosts transferred with TCR7B8 or TCR1A2 naïve CD4 T cells. Two independent experiments were conducted and the data from the indicated numbers of mice were pooled and reported as the mean ± SEM. **b,** Flow plots for TCRβ$^{+}$ cells from indicated tissues of TCR7B8 or TCR1A2 naïve CD4 T cell-transferred *Rag2*$^{-/-}$ hosts at 8 weeks following the transfer. Live MHCII$^{-}$ TCRβ$^{+}$ cells were highlighted with a circle gate, and the corresponding frequencies are as indicated. **c,** Numbers of IFNγ/IL-17A DP CD4 T cells in the indicated tissues 8 or 14 weeks following transfer. Two independent experiments were conducted and the data from the indicated numbers of mice were pooled and reported as the mean ± SEM. Student's t-test. ****P<0.0001, ***P<0.001, N.S.; not significant (P>0.05).

### Extended Data Figure 9

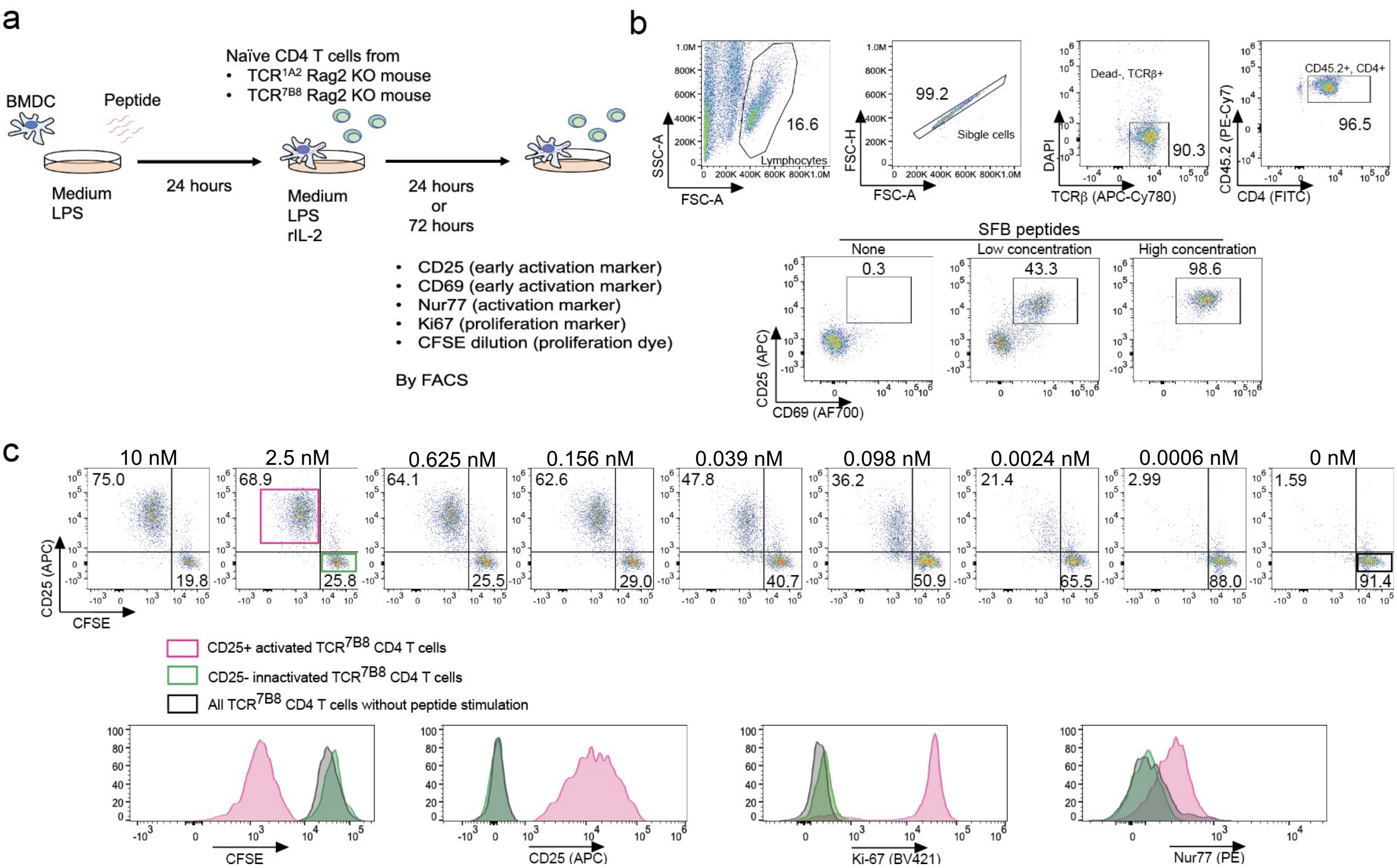


**Extended Data Fig. 9. In vitro stimulation assay using SFB-specific TCR$^{Tg}$ naïve CD4 T cells, BMDCs, and synthesized peptides.** Schematic representation of an *in vitro* naïve TCR$^{7B8}$ T cell stimulation assay using BMDCs and synthetic peptides. Synthesized peptides were mixed with 2-4x10$^4$ BMDCs induced from the BM of CD45.1/CD45.1 C57BL/6 mouse together with LPS in 96-well round-bottom plates. After 24 h, equal numbers of magnetically sorted naïve CD4 T cells from TCR$^{7B8}$ or TCR$^{1A2}$ animals on the CD45.2/CD45.2 *Rag2*$^{-/-}$ background were added together with recombinant IL-2. After an additional 24 h, all cells were collected and stained with anti-CD25, anti-CD69, anti-CD45.2, anti-TCRβ, and DAPI. For 72 h culture experiments, naïve CD4 T cells were labeled with CFSE just prior to co-culture with BMDCs. After 72 h, cells were stained with anti-CD25, anti-CD69, anti-CD45.2, anti-TCRβ, and Aqua. After fixation, cells were additionally stained with anti-Nur77 and anti-Ki67, and were then analyzed via FACS. **b,** The approach used to gate on CD25/CD69 double-positive proliferating TCR$^{Tg}$ CD4 T cells at 24 h following co-culture. **c,** Other proliferation and activation makers expressed on CD4 T cells were analyzed at 72 h following co-culture. CFSE-high CD25-negative non-proliferating T cells are highlighted with a black rectangle in non-peptide treated wells and with a green rectangle in the 2.5 nM peptide well. Effectively proliferating cells were defined as CD25+ cells exhibiting CFSE dilution, and these cells were gated with a red square and further assessed for Ki67 and Nur77 expression (bottom).

### Extended Data Figure 10

a

| ID | Peptide sequence | Gene name | ID | Peptide sequence | Gene name |
|---|---|---|---|---|---|
| 1 | water | | 40 | lqsesgavpkrk | spastin isoform 1 |
| 2 | vqfsgavpnktd | mouse SFB_003340 | 41 | eqvvlgqvpnkqk | rho guanine nucleotide exchange factor 5 |
| 3 | pvesgavprkpl | P2X purinoceptor 6 | 42 | pwaggdvphkth | dual specificity tyrosine-phosphorylation-regulated kinase 1B isoform p75 |
| 4 | srtsgavppkee | anaphase-promoting complex subunit 2 | 43 | gklsgatpngea | liprin-beta-2 isoform 1 |
| 5 | ltpsgavpnqaq | receptor tyrosine-protein kinase erbB-2 | 44 | ttksgqvpnlvt | neuronal-specific septin-3 isoform 11 |
| 6 | ppfsaalpnktm | RIKEN cDNA 1500035H01, isoform CRA_a, partial | 45 | yggsgnvpnrag | ankyrin repeat and SOCS box protein 15 |
| 7 | acgsgavpwklh | Ig-like V-type domain-containing protein FAM187A precursor | 46 | aqgvsgavqdkgs | inosine-5'-monophosphate dehydrogenase 2 isoform 2 |
| 8 | lafsgavqtkml | bile salt export pump | 47 | slssgaspyktc | pleckstrin homology-like domain family B member 2 isoform 2 |
| 9 | gqkfrgagpnkkv | spermatid perinuclear RNA-binding protein isoform X1 | 48 | pmatgavpakkk | alpha-N-acetylgalactosaminide alpha-2,6-sialyltransferase 1 |
| 10 | llfseavpikkr | epiplakin | 49 | wcgsgtvpgkqt | platelet-derived growth factor C isoform 1 precursor |
| 11 | tvfsgavstttg | trophinin isoform 1 | 50 | vhhsgavrnynr | speedy protein A isoform 1 |
| 12 | ilssgevpnlyk | dynein heavy chain 2, axonemal | 51 | eepsgpvpgkcp | receptor-interacting serine/threonine-protein kinase 3 isoform 1 |
| 13 | lefsgpipnpee | dishevelled associated activator of morphogenesis 2 | 52 | keksgavrnrkq | attractin preproprotein |
| 14 | rqfsgalplaqq | centrosomal protein of 290 kDa | 53 | ssnsgaspnpih | trinucleotide repeat-containing gene 6B protein isoform 1 |
| 15 | enpsnavpektq | 28S ribosomal protein S15, mitochondrial precursor | 54 | sprggavprkpv | ephrin-B3 precursor |
| 16 | gsligavpnstr | polycomb group RING finger protein 6 isoform X4 | 55 | gilsgatankas | probable hydrolase PNKD isoform 3 |
| 17 | dsfsgtvpaltl | uncharacterized protein CCDC198 isoform X1 | 56 | vsclgavvnkvt | nipped-B-like protein isoform a |
| 18 | gqtftssvpnker | dnaJ homolog subfamily C member 12 isoform 1 | 57 | prvsgavpgard | kelch domain-containing protein 3 |
| 19 | ssfsaaiptkts | TBC1 domain family member 5 isoform X1 | 58 | sptsgavppyss | myelin-associated neurite-outgrowth inhibitor isoform 3 |
| 20 | ryfsgalpdded | syndecan-4 precursor | 59 | lsgsgavpsmvv | membrane protein FAM174A precursor |
| 21 | rvfrgavpdrrr | mCG1045469 | 60 | scvsgatpnnst | mineralocorticoid receptor |
| 22 | pvfsgtvpgtpy | aggrecan core protein isoform X2 | 61 | kialgavpykee | centromere-associated protein E |
| 23 | hnfsgpvpesey | glycerol-3-phosphate acyltransferase 1, mitochondrial isoform X1 | 62 | yqvpgavpakvq | phospholipase D1 isoform 1 |
| 24 | ihfsgaipwick | regulator of G-protein signaling 1, isoform CRA_a, partial | 63 | tlgsgaspntgf | ribosomal protein S6 kinase-like 1 isoform 1 |
| 25 | tvfagavpvlpa | epidermal growth factor receptor substrate 15-like 1 | 64 | llqsgaimnkcy | DNA polymerase zeta catalytic subunit |
| 26 | tfsgvvpqtpa | RNA-binding protein 20 isoform X1 | 65 | Aqkssgavtkkgd | leucine-rich repeat-containing protein 71 |
| 27 | mlfsgavndmaa | TBC1 domain family member 8B isoform X1 | 66 | gvksgfvpntvh | ras-related protein Rab-39A |
| 28 | enfsaavphhrc | solute carrier family 22 member 12 isoform X1 | 67 | nkssgaqpnlkv | probable helicase senataxin |
| 29 | grfpgavpggdr | cleavage and polyadenylation specificity factor subunit 6 isoform X1 | 68 | svvsgadrnkae | O-phosphoseryl-tRNA(Sec) selenium transferase isoform X1 |
| 30 | tefpgavpkgvv | lysosome-associated membrane glycoprotein 2 isoform X1 | 69 | ssfsskvpevte | microtubule-associated protein 1A isoform X1 |
| 31 | aifpgavpaarp | forkhead box protein E1 | 70 | ksysnsvpektt | sodium-coupled monocarboxylate transporter 2 isoform 2 |
| 32 | ehfsgavyegqf | MORN repeat-containing protein 2 | 71 | rvcsgegppktd | Cadherin-related neural recepter 4 |
| 33 | ynfsgaapgppp | proline-rich protein 12 | 72 | kqevngavpddls | PWWP domain-containing protein 2A isoform b |
| 34 | lpfsgavsaqgi | collagen alpha-1(VII) chain | 73 | lqkregpvpsktn | augurin precursor |
| 35 | aafsgavthtls | Clcnkb protein | 74 | anisgsvplehf | aquaporin-7 isoform 2 |
| 36 | dmfsgavfiqqa | sodium/glucose cotransporter 2 | 75 | eldseavnnkvp | serine beta-lactamase-like protein LACTB, mitochondrial precursor |
| 37 | klcsgpvpnttq | thrombospondin type-1 domain-containing protein 7A | 76 | sassgsvsqktp | mediator of RNA polymerase II transcription subunit 1 isoform 3 |
| 38 | agysgavpqaea | ubiquitin thioesterase OTU1 | 77 | gggsggtpnttl | potassium voltage-gated channel subfamily D member 1 precursor |
| 39 | lrrygavpngfi | protein Dok-7 isoform 1 | 78 | mnpsgsvpegep | sialoadhesin precursor |

b

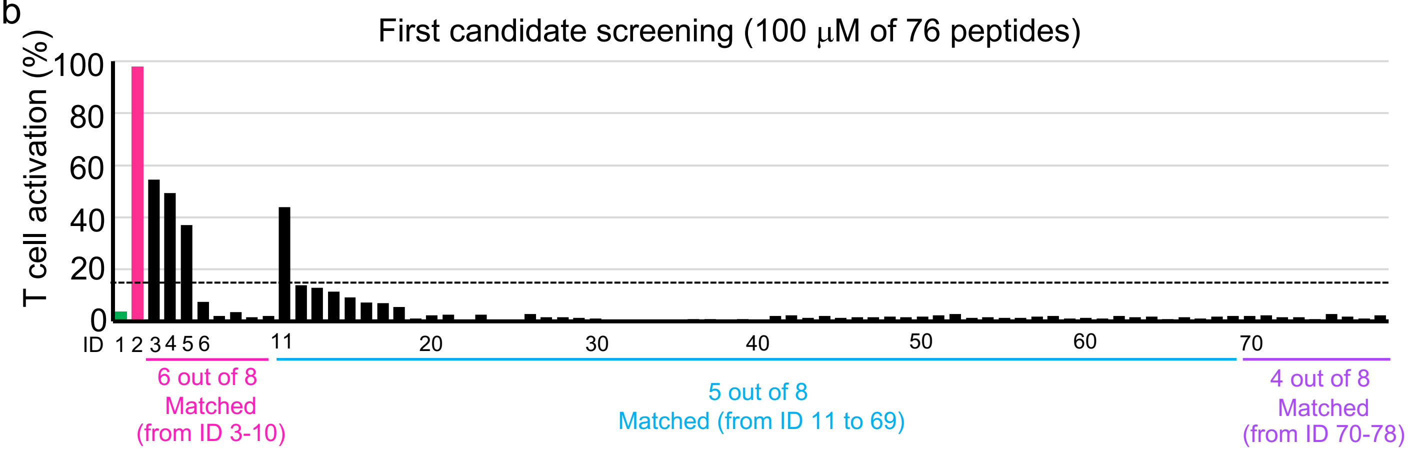


c

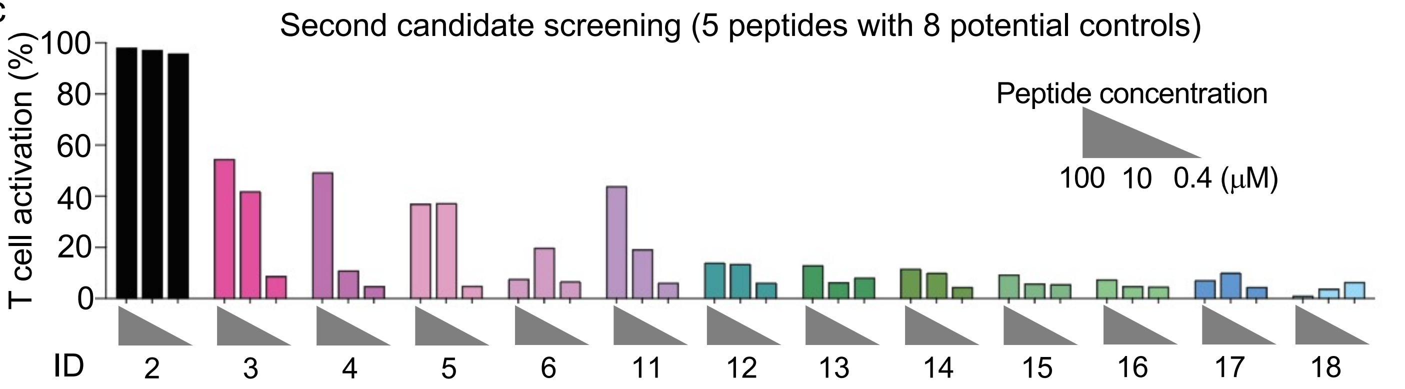

**Extended Data Fig. 10. Screening for potential cross-reactive candidate epitopes capable of stimulating naïve TCR$^{7B8}$ CD4 T cells *in vitro*.** **a**, A BLAST search was used to identify a list of candidate cross-reactive peptides. Core epitopes for TCR$^{7B8}$ and matched amino acids in candidate peptides are highlighted in red. Amino acids that did not match core epitopes for TCR$^{7B8}$ are shown in blue. **b**, The percentage of TCR$^{7B8}$ CD4 T cell activation in response to stimulation with a 100 μM concentration of candidate peptides. T cell activation was determined based upon CD25 and CD69 expression at 24 hours following co-culture. **c**, The dose-dependency of candidate peptide-induced T cell activation at 24 h following co-culture. In total, 11 candidates selected based on the results in (**b**) were further tested for their ability to activate TCR$^{7B8}$ CD4 T cells at concentrations of 10 μM and 0.4 μM. The results shown in (**b**) were also shown on the same graph in (**c**). Screening analyses were performed one time. P2X6 (ID3), ANAPC2 (ID4), ErbB2 (ID5), IFT46 (ID6), and TRO (ID11) were further generated as purified peptides and tested in Fig. 3.

### Extended Data Figure 11

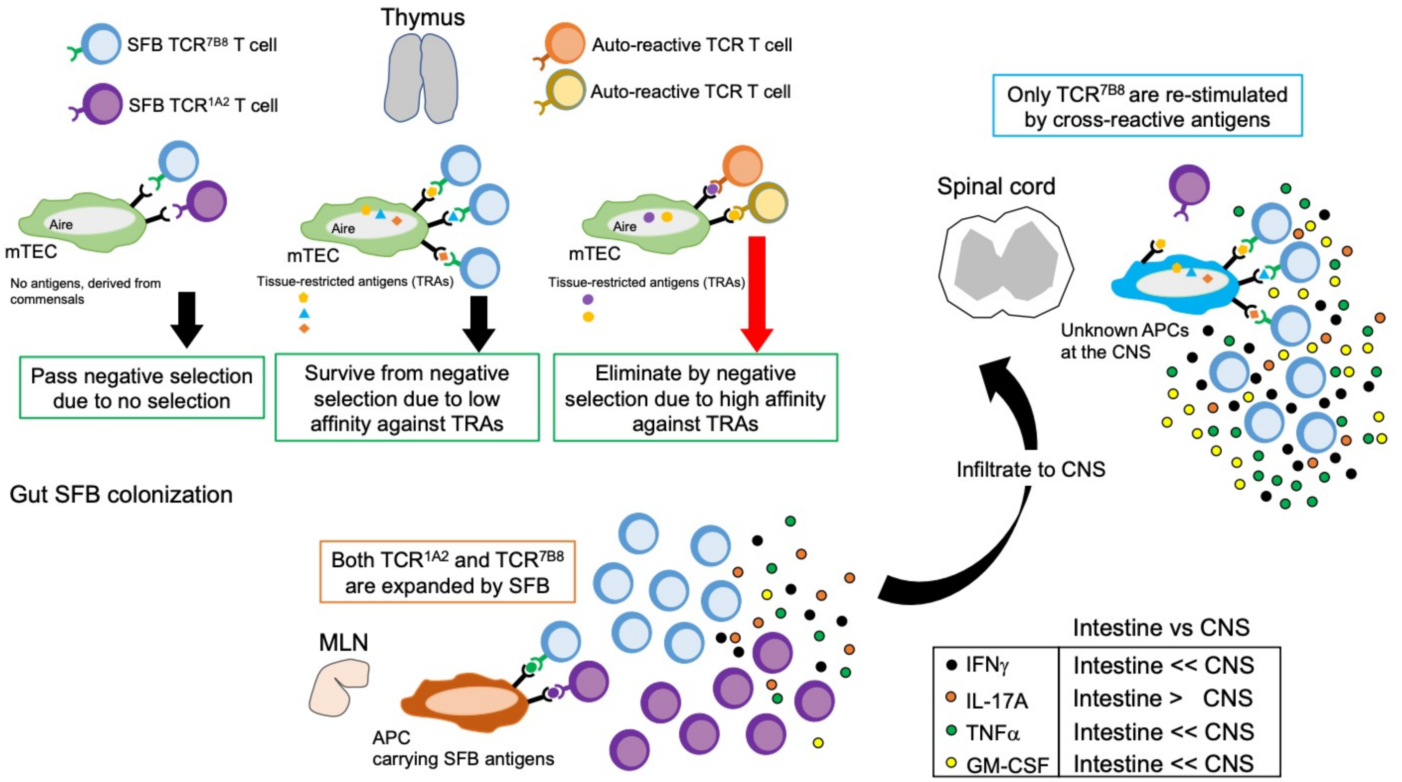


**Extended Data Fig. 11. Model for the mechanisms whereby TCR$^{7B8}$ T cells but not TCR$^{1A2}$ T cells contribute to CNS inflammation.** Commensal-specific T cells can be generated in the thymus and avoid negative selection. While TCR$^{7B8}$ T cells can cross-react with host protein-derived peptides, they likely escape negative selection owing to their low affinity for tissue-restricted antigens (TRAs) presented by mTECs. These commensal-specific naïve T cells can then travel to mLNs wherein they can differentiate into a range of helper T cell lineages upon commensal colonization. These differentiated commensal-specific T cells subsequently migrate to areas where cognate antigens are abundant, most often to the intestines although others can migrate to systemic sites such as the spleen. In the absence of Treg cells, SFB-specific TCR$^{7B8}$ and TCR$^{1A2}$ CD4 T cells are further activated in the small and large intestine and produce a range of effector cytokines, whereas Tregs suppress these activities. TCR$^{7B8}$ CD4 T cells can be found in CNS compartments including the brain and spinal cord in the absence of Tregs, and these tissue-infiltrating TCR$^{7B8}$ CD4 T cells produce higher levels of inflammatory cytokines such as TNF$\alpha$, GM-CSF, IFN$\gamma$, and IL-17A relative to cells found in the intestines, thus contributing to CNS inflammation.
